# Temperature effects on the magnitude of the bovine *ex vivo* immune response towards *Mycoplasmopsis bovis*

**DOI:** 10.1101/2023.10.16.562468

**Authors:** Thomas Démoulins, Thatcha Yimthin, Dorothea Lindtke, Lukas Eggerschwiler, Raphael Siegenthaler, Fabien Labroussaa, Joerg Jores

## Abstract

Although cattle are the mammalian species with most global biomass associated with a huge impact on our planet, their immune system remains poorly understood. Notably, the bovine immune system has peculiarities such as an overrepresentation of γδ T cells that requires particular attention, specifically in an infectious context. In line of 3R principles, we developed an *ex vivo* platform to dissect host-pathogen interactions. The experimental design was based on two independent complementary readouts: firstly, a novel 12-14 color multiparameter flow cytometry assay measuring maturation (modulation of cell surface marker expression) and activation (intracellular cytokine detection) of monocytes, conventional and plasmacytoid dendritic cells, natural killer cells, γδ T cells, B and T cells; secondly, a multiplex immunoassay monitoring bovine chemokine and cytokine secretion levels. The experiments were conducted on fresh primary bovine blood cells exposed to *Mycoplasmopsis bovis* (*M. bovis*), a major bovine respiratory pathogen. Besides reaffirming the tight cooperation of the different primary blood cells, we also identified novel key players such as strong IFN-γ secreting NK cells, whose role was so far largely overlooked. Additionally, we compared the host-pathogen interactions at different temperatures, including commonly used 37 °C, ruminant body temperature (38-38.5 °C) and fever (≥39.5 °C). Strikingly, working under ruminant physiological temperature influenced the capacity of most immune cell subsets to respond to *M. bovis* compared to 37 °C. Under fever-like temperature conditions the immune response was impaired compared to physiological temperature. Our experimental approach, phenotypically delineating the bovine immune system provided a thorough vision of the immune response towards *M. bovis* and the influence of temperature towards that immune response.

## INTRODUCTION

Bacteria of the class *Mollicutes* encompass the smallest and simplest self-replicating bacteria. They are characterized by the absence of cell wall and a small, streamlined genome, as a result of reductive evolution. Among them, *Mycoplasmopsis bovis* (*M. bovis*) is one of the major causative pathogens of bovine respiratory complex disease (BRD), characterized among others by enzootic pneumonia, pleuritis and polyarthritis [1, 2]. The lack of cell wall has two major consequences. Firstly, lipid and lipoprotein antigens are anchored on the outer face of the plasma membrane, thus leading to high exposure to the host. Consequently, these antigenic structures are likely to elicit strong host immune responses, generally of inflammatory nature, albeit also being able to evade the immune system, requiring a better understanding (for a review, see [3]). Secondly, the lack a cell wall renders *M. bovis* inherently resistant to a large spectrum of antibiotics; effectively, available antibiotics such as tetracycline and spectinomycin are most of time ineffective [4, 5], pointing out the necessity to develop novel vaccines to circumvent the antibiotic treatment.

Currently, only a limited number of vaccines are available against ruminant *Mollicutes* and many have shortcomings, impacting their acceptance and use in the field [6]. The development of rationally designed vaccines benefits from a good understanding of host-pathogen interactions. Although cattle are the mammalian species with most planetary biomass associated with a huge impact on our planet [7], immune cell subset research in cattle is relatively young and remains largely elusive due to the lack of commercial reagents. Despite this limitation, striking differences were found between bovine and human monocyte subsets regarding their *in vitro* function [8, 9]. Another major distinction is the overrepresentation of γδ T cells in cattle, although still incompletely understood [10]. γδ T cells express transmembrane glycoproteins called workshop cluster 1 (WC1), also involved in the pattern recognition of bacteria [11]. Recently, some efforts have been made to better characterize mainly subsets from innate immunity or cells bridging innate-adaptive immunity: monocyte (classical, intermediate and non-classical) and blood dendritic cells (conventional DC type 1 and 2 (cDC1, cDC2), plasmacytoid DC (pDC)) and monocytes [12–14]. This, added to the fact *M. bovis* is known for decades to modulate bovine immune responses [15, 16], offers an interesting frame for reassessing host-*M. bovis* interaction studies.

In line with the 3R principles, we developed an *ex vivo* laboratory platform employing peripheral blood mononuclear cells (PBMCs) of outbred animals to investigate bovine-pathogen interactions. The first aim was to setup an assay allowing the precise delineation to other subsets than monocytes and DCs: cell types from adaptive immunity (B and T cells), as well as cell types linking the innate and adaptive arms of the immune system (Natural killer (NK) and γδ T cells)). Therefore, additional markers were included in the panel to quantify with accuracy the activation and maturation state of 13 investigated cell subsets. This, coupled with multiplex immunoassay, commercially available only since 2019, allowed us also to monitor chemokine and cytokine secretion, and provided a powerful experimental set-up for an unprecedented deciphering of concerted actions of bovine immune cells using an *ex vivo* format. We tested the assay using *M. bovis,* an important bacterial pathogen, causing arthritis, mastitis and pneumonia in cattle. Finally, the present study gave new insights on the consequences of elevated body temperature on the immune responses against bacterial pathogens, a so far largely overlooked parameter. Intriguingly, we found that high fever temperature negatively impacted the capacity of PBMCs to respond to *M. bovis*.

## MATERIALS AND METHODS

### Mycoplasmopsis bovis Donetta PG45 strain

*M. bovis* Donetta PG45 strain was obtained from the American Type Culture Collection (ATCC, 25523). *M. bovis* was grown statically in tubes at 37 °C and 5 % CO_2_ in Mycoplasma Experience Liquid medium (Mycoplasma Experience, Bletchingley, Surrey, UK) overnight, then harvested after changed color by pelleting the culture at 3,400 x g for 15 min. *M. bovis* were re-suspended in fresh Mycoplasma Experience medium, aliquoted in 1 ml and stored at −80 °C until further use.

To determine the actual *M. bovis* titer, an aliquot was thawed, spot diluted on Mycoplasma Experience agar plates (Mycoplasma Experience) and incubated at 37 °C and 5 % CO_2_ for 5 days before determining the colony forming units (CFU).

To heat-inactivate *M. bovis*, an aliquot was thawed and heated at 65 °C for 10 minutes before being used for primary blood cell stimulation.

### Isolation of bovine peripheral blood mononuclear cells

Blood of Holstein Friesian cows (aged 1–3 years) was collected at the Agroscope research facility (Posieux, Switzerland) for the main assays (n=8 animals) (**Additional file 1**), or at the Institute of Virology and Immunology (IVI, Mittelhäusern, Switzerland) for the preliminary experiments (n=2 animals). 50-100 ml of blood was obtained by puncturing the jugular vein into vacutainer EDTA tubes (Becton Dickinson). For peripheral blood mononuclear cell (PBMC) isolation, blood was first centrifuged at 1,000 x g (20 min). Then the buffy coat was collected and diluted with PBS containing 1 mM UltraPure™ EDTA (Invitrogen, ThermoFisher) to a ratio of 1 to 1 before being layered onto Ficoll Paque (1.077 g/ml; GE Healthcare Europe GmbH, Freiburg, Germany). After centrifugation (800 x g for 25 min), PBMCs were collected and washed twice with cold PBS containing 1 mM EDTA (Invitrogen) (350 x g for 10 min). A final washing step was done at 250 x g for 10 min to remove platelets.

The donor cattle enrolled in the assays were from acknowledged animal facilities in Switzerland and represent herds that are not mingling with other animals outside the facilities. Experimental animals enrolled in this study had no previous reports of *M. bovis*-related disease. Moreover, lack of previous exposure towards *M. bovis* was unequivocal confirmed on all animal serums via ID Screen Mycoplasma bovis indirect ELISA (Innovative Diagnostic, France). Effectively, results in **Additional file 2** clearly showed that all samples had an optical density by far < 60% of that quantified in positive control, threshold demonstrating no previous contact with *M. bovis* that could affect our results and the subsequent immunological interpretations.

### Whole genome sequencing and kinship analysis of cattle used in this study

Genomic DNA was extracted from EDTA-blood of 21 cows, employing the Maxwell^®^ RSC Whole Blood DNA Kit and the Maxwell^®^ RSC Instrument, and sequenced on an Illumina NovaSeq 6000 using 150 bp paired-end sequencing libraries. Raw reads were trimmed and filtered with the fastp software [17] version 0.23.2 by enabling adapter sequence auto-detection, removing one base at the front of each read, additionally trimming one base at the front or tail of the read if it had a Phred quality below 5, and trimming the read if the mean Phred quality in a sliding window of four bases (moving from front to tail) was below 20. Reads shorter than 30 bp after trimming were discarded, and base correction in overlapping regions between read pairs was enabled. Default settings were used for all other parameters. Filtered paired reads were mapped separately by sample, flow cell and lane to the bovine reference genome version ARS-UCD1.3 (GCF_002263795.2) using the mem algorithm implemented in bwa-mem2 [18] version 2.2.1. Picard tools [https://broadinstitute.github.io/picard] version 3.0.0 was employed to merge and coordinate-sort the resulting SAM files for each sample and to mark duplicate reads.

GATK version 4.4.0.0 [19] was used to recalibrate base quality scores, based on known *Bos taurus* variants from Ensembl release 109, and to discover polymorphic sites using the Best Practices Workflow [20] under default settings. Biallelic SNPs were extracted and hard-filtered according to the GATK recommendations (i.e., removing SNPs with QD < 2.0, FS > 60.0, SOR > 3.0, MQ < 40.0, MQRankSum < -12.5, or ReadPosRankSum < -8.0) and additionally removing SNPs with read depth (DP) < 21 or DP > mean(DP) + 5 * sd(DP). SNPs with a minor allele frequency below 0.05 or any missing genotypes were discarded using VCFtools [21] version 0.1.17.

The filtered VCF file was converted to BED format using PLINK2 [22] version 2.0.0a3.3, and kinship was estimated from autosomal SNPs using the robust algorithm of KING [23] version 2.3.0. Autosomal SNPs were thinned to contain at most one random variant per 200 bp using PLINK2, and used to estimate inbreeding coefficients based upon the correlation between uniting gametes and to perform principal component analysis (PCA) on the genetic relationship matrix with GCTA [24, 25] version 1.94.1.

### Stimulation of bovine peripheral blood mononuclear cells

Blood cell stimulation employed flat-bottom 6-well plates (TPP, Switzerland). Per well, 5 × 10^6^ cells were cultured in 3 ml Dulbecco’s modified Eagle’s medium (DMEM) (Life Technologies) supplemented with 10% fetal bovine serum (Life Technologies).

The synthetic TLR ligands Pam3Cys-SK4 (10 μg/ml), polyinosine–polycytodylic acid (polyI:C) (10 μg/ml), lipopolysaccharide (LPS) (10 μg/ml) and resiquimod (R848) (10 μg/ml), all from InvivoGen, were used for stimulation of cells over 18 h. Likewise the mitogens phorbol 12-myristate 13-acetate (PMA) (200 ng/ml), ionomycin (1 μg/ml) and concanavalin A (10 μg/ml), all from Sigma-Aldrich, were used for stimulation of cells over 18 h.

Stimulation with *M. bovis* (live or heat-inactivated) was done at a multiplicity of infection (MOI) of 0.1, based on preliminary experiments, for 18 h.

After 14 h, 50-100 μl of cell culture supernatant was collected and frozen for further cytokine secretion measurement (multiplex immunoassay). As soon after, Brefeldin A (10 μg/ml) (ThermoFisher) was added to the medium to block cytokine secretion; incubation was extended for another 4 h, to allow the de novo cytokine synthesis measurement (flow cytometry intracellular staining).

### Multiparameter flow cytometry assay

Counting of *M. bovis* by flow cytometry (FCM) employed *Bac*Light^TM^ RedoxSensor^TM^ Green Vitality Kit, following manufacturer’s instructions (ThermoFisher).

The different bovine immune cell subtypes were identified by FCM using a 7-step, 11-12-color staining protocols, based on recent reports [26, 27]. All antibodies used for the procedure, as well as their clones, host, working dilution, and references, are listed in **Table 1**. Combination stainings analyzed monocytes (conventional, intermediate and non-conventional), cDCs (cDC1s and cDC2s), pDCs, γδ T cells, NK cells, CD4^+^ and CD8^+^ T cells, and B cells. For the acquisitions, at least 10^6^ events were accumulated for each sample. The experimental design is depicted in **Fig. 1**.

**Figure 1.**
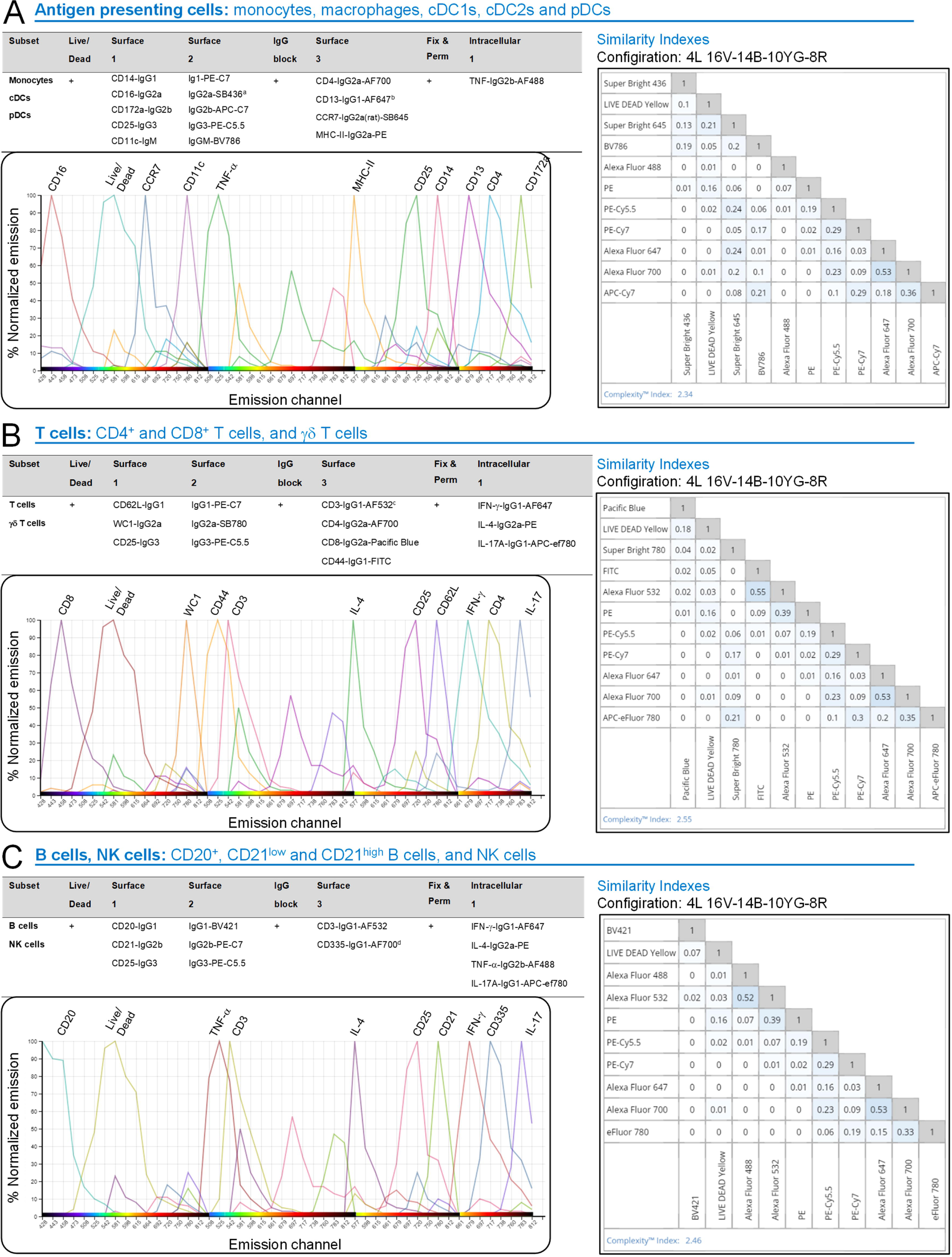
Combinatorial immunostainings used for the identification of bovine immune cell subsets. The staining procedure used a seven-step, eleven-color assay. Three panels were designed for in depth characterization of antigen presenting cells (i.e., monocytes, macrophages and DCs) **(A)**, T cells (CD4^+^, CD8^+^ and γδ T cells) **(B)** and B and NK cells **(C)**. The Similarity Index is a measure of dye pair uniqueness on a scale from 0 (the 2 dyes are very divergent from each other) to 1 (the 2 dyes are very similar to each other), indicating that none of our dyes were incompatible with each other (acceptable threshold ≤ 0.98). The Complexity Index is an overall measure of uniqueness of all dyes in a full spectrum cytometry panel. Our three eleven-color combinations have a complexity Index < 3, proving they were well-designed panels according to the manufacturer. Emission channel Illustrations are taken from Cytek Full Spectrum Viewer website. ^a^ Due to product discontinuation, the antibody combination panel was occasionally slightly modified: IgG1-AF405 and IgG2a-PE-C7 rather than IgG1-PE-C7 and IgG2a-SB436, respectively. ^b^ Labeled with Zenon™ Mouse IgG1 Labeling Kit, AF647. ^c^ Labeled with AF532 Antibody Labeling Kit. ^d^ Labeled with Zenon™ Mouse IgG1 Labeling Kit, AF700.

**Table 1.** Antibodies used in the study.

| Antibody | Source | Identifier |
| --- | --- | --- |
| Anti-CD14 (Clone CAM36A; mouse Ig1; 1:800 dilution) | Kingfisher Biotech | Cat# WS0565B-100 |
| Anti-CD25 (Clone LCTB2A; mouse IgG3; 1:200 dilution) | Kingfisher Biotech | Cat# WS0597B-100 |
| Anti-CD11c (Clone BAQ153A; mouse IgM; 1:40 dilution) | Kingfisher Biotech | Cat# WS0519B-100 |
| Anti-CD16 (Clone KD1; mouse, IgG2a; 1:100 dilution) | Bio-Rad | Cat# MCA5665GA |
| Anti-CD172a (Clone CC149; mouse IgG2b; 1:200 dilution) | Bio-Rad | Cat# MCA2041GA |
| Anti-CD62L (Clone CC32; mouse IgG1; 1:320 dilution) | Bio-Rad | Cat# MCA1649G |
| Anti-WC1 (Clone CC15; mouse IgG2a; 1:800 dilution) | Bio-Rad | Cat# MCA838 |
| Anti-CD21 (Clone CC51; mouse IgG2b; 1:800 dilution) | Bio-Rad | Cat# MCA5953 |
| Anti-CD4-AF700 (Clone CC8; mouse IgG2a; 1:200 dilution) | Bio-Rad | Cat# MCA1653A700 |
| Anti-CD13 (Clone CC81; mouse IgG1; 1:100 dilution) | Bio-Rad | Cat# MCA2338GA |
| Anti-MHC-II-PE (Clone IL-A21; mouse IgG2a; 1:40 dilution) | Bio-Rad | Cat# MCA2445PE |
| Anti-CD3 (Clone MM1A; mouse IgG1; 1:300 dilution) | Bio-Rad | Cat# MCA6080 |
| Anti-CD8-Pacific Blue (Clone CC63; mouse IgG2a; 1:20 dilution) | Bio-Rad | Cat# MCA837PB |
| Anti-CD44-FITC (Clone IL-A118; mouse IgG1; 1:40 dilution) | Bio-Rad | Cat# MCA2433F |
| Anti-CD335 (Clone AKS1; mouse IgG1; 1:100 dilution) | Bio-Rad | Cat# MCA2365GA |
| Anti-TNF-α-AF488 (Clone CC327; mouse IgG2b; 1:80 dilution) | Bio-Rad | Cat# MCA2334A488 |
| Anti-IFN-γ-A647 (Clone CC302; mouse IgG1; 1:160 dilution) | Bio-Rad | Cat# MCA1783A647 |
| Anti-IL-4-PE (Clone CC303; mouse IgG2a; 1:20 dilution) | Bio-Rad | Cat# MCA1820PE |
| Anti-CD20 (Clone MEM-97; mouse IgG1; 1:100 dilution) | TermoFisher | Cat# MA1-19008 |
| Anti-CCR7-Super Bright 645 (Clone 3D12; Rat IgG2a; 1:80 dilution) | TermoFisher | Cat# 64-1979-42 |
| Anti-IL-17A-APC-ef780 (Clone Ebio64DEC17; mouse IgG1; 1:160 dilution) | TermoFisher | Cat# 47-7179-42 |
| Anti-mouse IgG2a-Super Bright 436 (Clone m2a-15F8; rat IgG1; 1:20 dilution) | TermoFisher | Cat# 62-4210-82 |
| Anti-mouse IgG2a-Super Bright 780 (Clone m2a-15F8; rat IgG1; 1:40 dilution) | TermoFisher | Cat# 78-4210-82 |
| Anti-mouse IgG3-PE-Cy5 (Goat IgG; 1:200 dilution) | SouthernBiotech | Cat# 1100-13 |
| Anti-mouse IgG2b-APC-Cy7 (Goat IgG; 1:400 dilution) | SouthernBiotech | Cat# 1090-19 |
| Anti-mouse IgG2b-PE-Cy7 (Goat IgG; 1:250 dilution) | SouthernBiotech | Cat# 1090-17 |
| Anti-mouse IgG1-PE-Cy7 (Clone RMG1-1; rat IgG; 1:640 dilution) | BioLegend | Cat# 406614 |
| Anti-mouse IgG1-Brilliant Violet 421 (Clone RMG1-1; rat IgG; 1:80 dilution) | BioLegend | Cat# 406616 |
| Anti-mouse IgM-BV786 (Clone R6-60.2; rat IgG2a; 1:100 dilution) | BD Bioscience | Cat# 564028 |
| Others | Source | Identifier |
| Zenon™ Mouse IgG1 Labeling Kit, AF647 | TermoFisher | Cat# Z25008 |
| AF532 Antibody Labeling Kit | TermoFisher | Cat# A20182 |
| Zenon™ Mouse IgG1 Labeling Kit, AF700 | TermoFisher | Cat# Z25011 |
| LIVE/DEAD™ Fixable Yellow Dead Cell Stain Kit, for 405 nm | TermoFisher | Cat# L34968 |
| Brefeldin A Solution (1000X) | TermoFisher | Cat# 00-4506-51 |

For the fold-change analysis of activation / maturation marker following stimulation, the mean fluorescence intensity (MFI) measured in stimulated sample for a given cow was normalized to the MFI measured in unstimulated sample from that same animal.

FCM acquisitions were performed on a Cytek Aurora (Cytek Biosciences) using the SpectroFlo software with autofluorescence extraction, and further analyzed with FlowJo 10.8.1 (TreeStar).

### Multiplex immunoassay

Bovine cytokines and chemokines secreted in the supernatant by cells were measured using the commercial multiplex immunoassay MILLIPLEX MAP Bovine Cytokine/Chemokine Panel 1 (Merck/Sigma). Therefore, supernatants of cultured PBMCs were collected, aliquoted and frozen a few months at -20 °C until further use. For analysis, the supernatants were thawed, and cytokine/chemokine concentrations were assessed using the immunoassay according to the manufacturer’s protocol. Samples were acquired on a Luminex™ xMAP™ INTELLIFLEX including an xPONENT Software version 4.2 software (Luminex) and further analyzed with Belysa™ software.

### Statistical analysis

Statistical analysis was done using the GraphPad Prism 8 software (GraphPad software, La Jolla, CA, USA). To determine differences between groups, paired t tests or one-way repeated measure ANOVA followed by Tukey’s post hoc test were used, as appropriate. Associations were tested using the Spearman rank correlation test. A p value < 0.05 was considered statistically significant (* p < 0.05, ** p < 0.01, *** p < 0.001, **** p < 0.0001).

## RESULTS

### Determination of the kinship relationship of cows enrolled in the study

Whole genome sequencing of 21 female Holstein cows resulted in an average of 234 M raw read pairs per sample (range: 215 M to 297 M), of which on average 99.2% were retained after filtering. The average read mapping rate against the reference genome was 99.4% and resulted in an average depth of 23.6x (range: 21.7x to 30.0x). A total of 15.6 M raw SNPs and 2.5 M raw INDELs were called in the joint set, of which 10.7 M biallelic SNPs passed filtering. Of those, 10.2 M were located on autosomes, and 4.3 M remained after thinning.

Kinship estimates revealed the presence of some 2nd and 3rd order relationships between animals analyzed, including one cluster of seven cows (**Additional file 3**), which explained 10.28% of the genetic variance among all animals (**Additional file 4**). Inbreeding coefficients were on average -0.03 (range: - 0.10 to 0.04). Consequently, the animals enrolled in the present study can be considered as largely outbred and genetic heterogeneous, supporting the external reproducibility of our findings (**Additional file 3** and **Additional file 4**) [28].

### Combinatorial immunostainings used for the identification of bovine immune cell subsets enable the setup of the FCM-based readout assay

We aimed to establish an FCM-based readout to delineate bovine immune components, which required an in-depth literature study including the availability of specific antibodies for cattle. After identification of available biologicals, we run a number of preliminary experiments to determine (i) their individual working dilution (to reduce cost and undesired background) and (ii) their compatibility when used in combination. All the antibodies selected, validated, and titrated in the study are listed in **Table 1**. As a result, three immunostaining combinations were designed with low similarity index (SI), indicative of low overlap – and thus good compatibility – of the different dyes: “Antigen presenting cells (APCs)”, encompassing monocytes, macrophages, cDC1s, cDC2s and pDCs (SI=2.43); “T cells”, encompassing CD4^+^ and CD8^+^ T cells, γδ T cells” (SI=2.49); and “B cells, NK cells”, encompassing CD20^+^, CD21^low^ and CD21^high^ B cells, as well as NK cells (SI=2.49). **Fig. 1** summarizes the three distinct immunostainings, detailing the strategy involving a 7-step, 10-11-color assay. Examples of gating strategy for the aforementioned immune cell subset identifications are provided in **Additional file 5.**

### Synthetic TLR ligands stimulate primary blood cells, which can be monitored by the assay

To ascertain the responsiveness of bovine PBMCs and the ability of our immunological platform to report it, blood cells from eight individual cattle were stimulated for 18 h, with either TLR2/4 (“bacterial infection”) and/or TLR3/7/8 (“viral infection”) agonists.

Our multiparameter FCM assay was first applied to evaluate the influence of TLR2/4 and TLR3/7/8 agonists on APC activation/maturation, via the quantification of MHC-II (related to antigen presentation), CD25 (related to cellular activation) and CCR7 (related to migration towards draining lymph node) expression. Interestingly, TLR2/4 stimulation influenced all monocyte subsets by dampening the MHC-II expression level (normalized values below 1); this trend was also observed for TLR3/7/8, but clearly to a lesser extent (**Fig. 2A**). This finding was not observed in DCs, the major APC subset. These cells have a very high MHC-II expression level under basal conditions, which do not ease the visualization of slight modulations. Nonetheless, when we considered the CD25 readout, the upregulation obtained following TLR2/4 stimulation was clearly further increased upon TLR3/7/8 stimulation for all DC subsets (**Fig. 2B**). Consequently, similarly to monocytes, DCs were differentially modulated by conditions mimicking “bacterial infection” or “viral infection”. Of note, TLR ligand exposure had an inconsistent influence on CCR7 expression level: in some subsets had TLR2/4 the stronger effect (on intermediate monocytes), whereas in other subsets was TLR3/7/8 more effective (cDC2s and pDCs). Therefore, not only could we induce a fine-tuning of the activation/maturation state of APC subsets, but we also proved that our new experimental tool quantified appropriately those differential changes.

**Figure 2.**
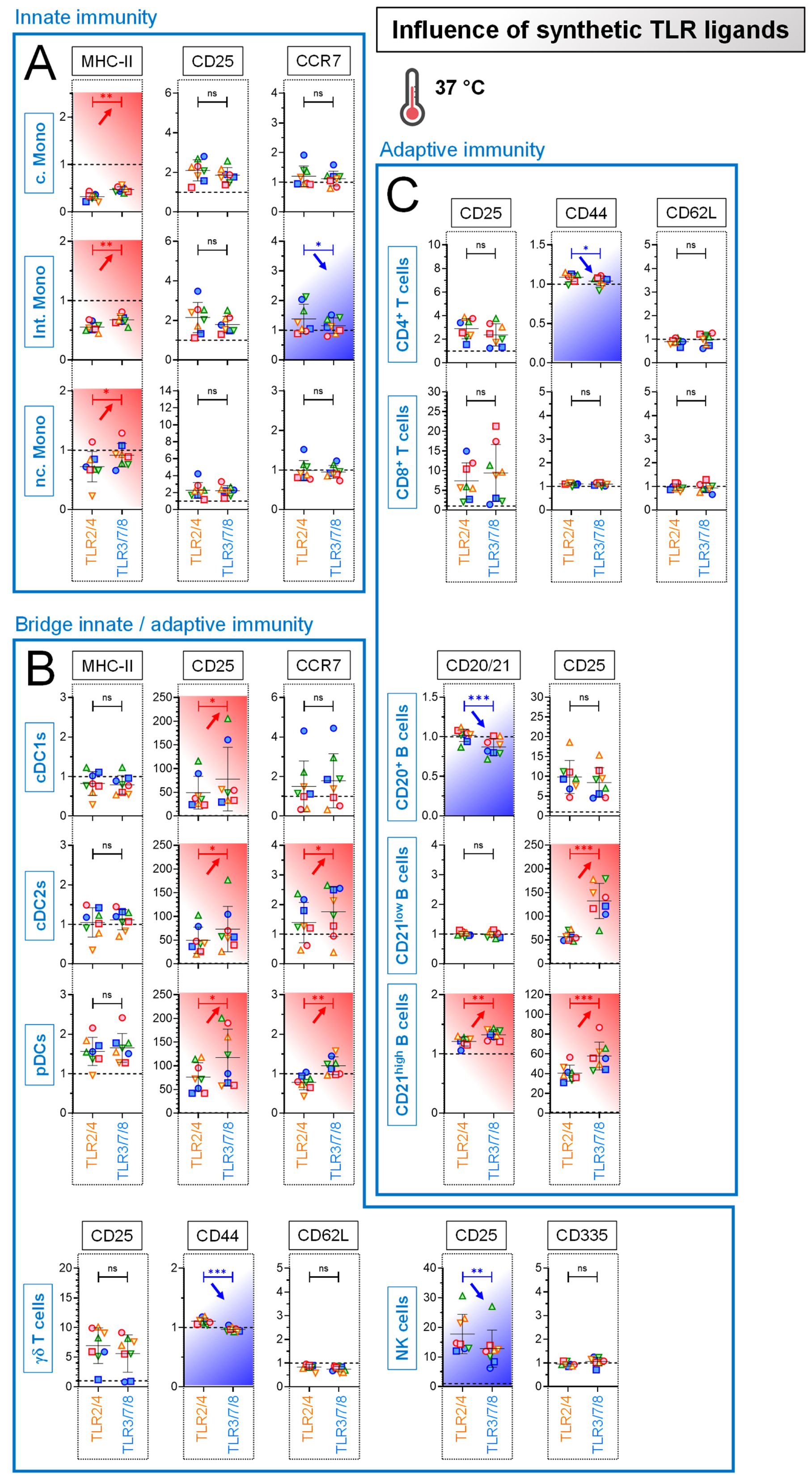
Stimulation of primary blood cells by synthetic TLR ligands. PBMCs from 8 individual animals were either let for 18 h unstimulated (reference points for the assay), stimulated with TLR2/4 (“bacterial infection”) or stimulated with TLR3/7/8 (“viral infection”). The fold changes analysis of activation / maturation markers was determined by FCM with FlowJo. Cells from the individual cows arerepresented by separate symbols; for stimulated samples, mean fluorescence intensity (MFI) values are normalized to that obtained with the reference point from the same animal. (**A**) Results obtained for the “Innate immunity” cell subsets. (**B**) Results obtained for the “Bridge Innate/adaptive immunity” cell subsets. (**C**) Results obtained for the “Adaptive immunity” cell subsets. Red arrows indicate a further increase of activation/maturation marker expression induced by TLR3/7/8 (“viral infection”) compared to TLR2/4 (“bacterial infection”); blue arrows indicate a decrease of activation/maturation marker expression. Experimental conditions were compared using paired t tests. Stars indicate significance levels. *, p < 0.05; **, p < 0.01; ***, p < 0.001; ns: not significant. c. Mono: classical monocytes; int. Mono: intermediate monocytes; n.c. Mono: non classical monocytes.

Our FCM assay was next applied to characterize the response of cells bridging innate-adaptive immunity (γδ T cells and NK cells), and cells from adaptive immunity (T and B cells). Again, TLR2/4 and TLR3/7/8 showed distinct modulatory capacity on most of those immune cell subsets (with the exception of CD8^+^ T cells, whose the differential modulation required stimulation by separate mitogens (data not shown)). The more active synthetic agonist could be either TLR2/4 (for example CD25 upregulation on NK cells) (**Fig. 2B**), or TLR3/7/8 (for example CD25 upregulation on CD21^+^ B cells) (**Fig. 2C**). In conclusion, depending on the “bacterial infection” or “viral infection” mimicking conditions, a differential immunomodulation of the vast majority of immune cell subsets (innate, adaptive, bridge innate-adaptive) could be measured. Collectively, these results showed that our newly designed experimental setup i) deciphers appropriately qualitative and magnitude changes in the activation/maturation state of a large panel of immune cell subsets, and ii) is a prelude to examine numerous host-live pathogen interactions.

### Cytokines are secreted by primary blood cells following exposure to synthetic TLR ligands

Next, we aimed to confirm the distinct immunomodulatory capacity of TLR2/4 and TLR3/7/8 on PBMCs with a different, FCM-independent technology, namely the Multiplex Immunoassay. As expected for any TLR triggering, both kind of stimulations induced particularly elevated amounts of pro-inflammatory cytokines (IL-1β, IL-6 and TNF-α), as well as high levels of chemokine MIP-1β. More surprising was the paralleled production of the anti-inflammatory cytokine IL-10 (**Fig. 3A** and **3B**).

**Figure 3.**
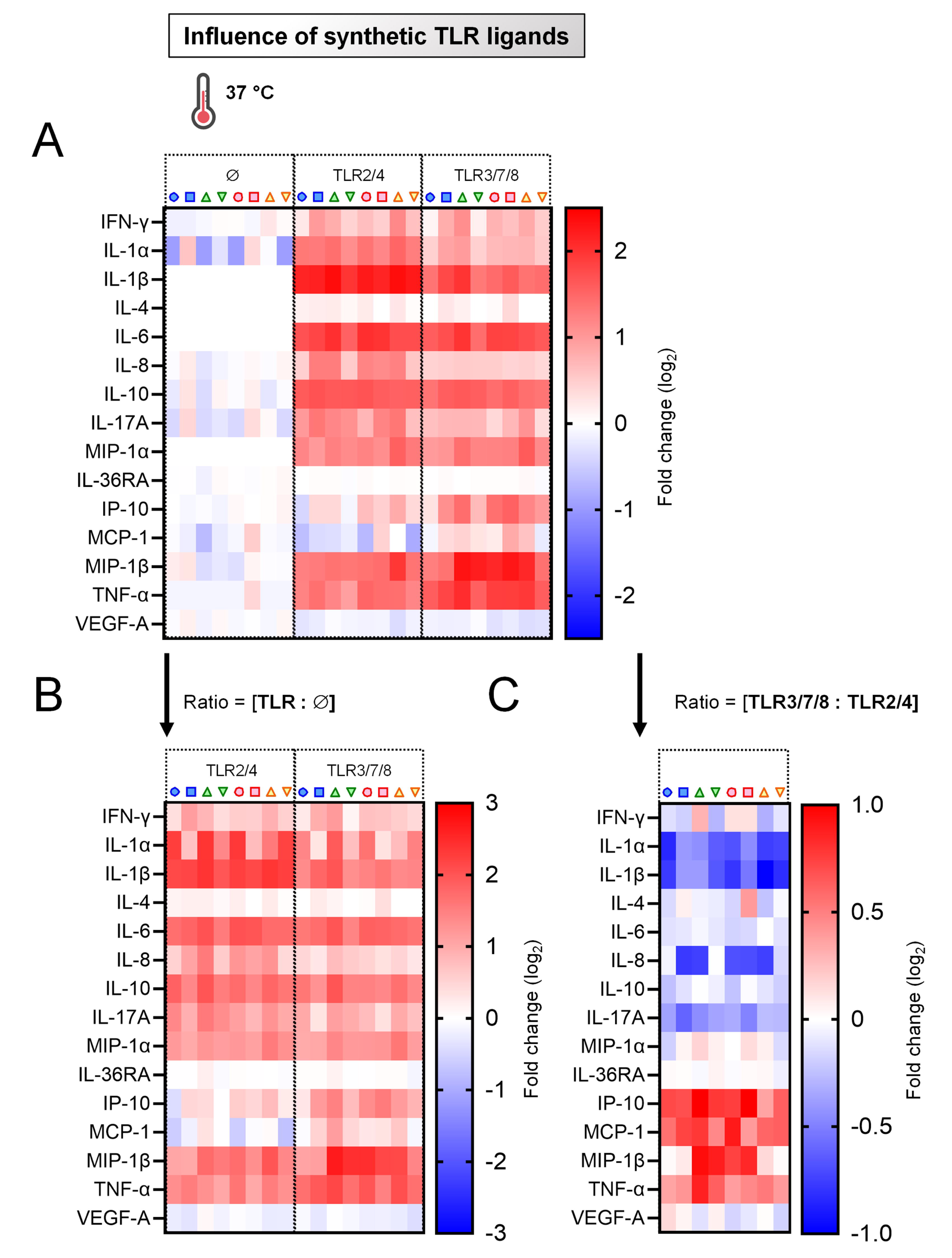
Induction of cytokines by primary blood cells following exposure to synthetic TLR ligands. PBMCs from 8 individual animals were either let for 18 h unstimulated (reference points for the assay), stimulated with TLR2/4 (“bacterial infection”) or stimulated with TLR3/7/8 (“viral infection”). (**A**) Cytokine/chemokine secretion in supernatants of PBMC cultures, using Multiplex immunoassay. A single measurement was done per samples tested, and each symbol represents an individual cow. Heat map shows log_2_-fold changes in concentration of 15 cytokines/chemokines. For a given cytokine / chemokine, normalization was as follow: [concentration for a given animal] / [average concentration of reference points Ø for 8 animals]. (**B**) As in (A), but displaying the ratios: [TLR2/4 : Ø] and [TLR3/7/8 : Ø]. (**C**) As in (A), but displaying the ratio: [TLR3/7/8 : TLR2/4].

When the values obtained with TLR3/7/8 were normalized to data obtained with TLR2/4, a divergent influence was noted whether we considered IL-1α, IL-1β, IL-17A (negative fold change, blue color patterns), or IP-10, MCP-1 and TNF-α (positive fold change, red color patterns) (**Fig. 3C**). Again, this showed i) the capacity of TLR2/4 and TLR3/7/8 combinations to differentially modulate *ex vivo* bovine primary blood cells, as well as ii) the robustness of our experimental approach combining multiparameter FCM and Multiplex immunoassays to study immune responses in cattle.

### Effect of cell culture medium on *Mycoplasmopsis bovis* fitness and viability

*M. bovis* is the etiological agent for many disorders in cattle, including mastitis, pneumonia and arthritis for which recently the commercial live-attenuated vaccine Protivity™ has been released. Also, a bacteremia has been reported in the course of *M. bovis* infection. Our immunological assays require to be performed in culture medium suitable for bovine PBMCs (**Additional file 6, A**). We thus tested whether *M. bovis* growth and/or survival were affected in DMEM supplemented with 10% FBS (DMEM/10%).

To this aim, *M. bovis* was grown and harvested at different time points in mycoplasma experience medium (MEX) or DMEM/10%, either at 37 °C (standard temperature for most immunological assays, below the bovine standard body temperature of 38.5 °C, “Hypothermia”) or 39.5 °C (“Low fever”), postulating that the later is more related to bacterial infection. When CFU was used as readout, the different culture conditions produced comparable CFUs up to 9 h after incubation, however with a trend to increased numbers upon incubation in MEX, at 39.5 °C (**Additional file 6, B**).

We then employed FCM to assess more precisely *M. bovis* growth in the different aforementioned culture conditions. Consistent with previous observations, the bacteria proliferated better in MEX (**Additional file 6**, C, upper panel). Next, we evaluated whether the reduction in viable *M. bovis* numbers observed in DMEM/10% was due to growth inhibition or death. Firstly, FCM acquisition showed a slight increase of *M. bovis* numbers in DMEM when incubation was extended from 6 h to later time point (18 h): this clearly indicated that *M. bovis* continued to grow. But this appeared to be minor compared to the *M. bovis* counts measured in MEX (18.2-fold increase). Secondly, a viability assay was conducted; it showed that incubation in DMEM/10% resulted in approximatively 5% of dead *M. bovis* cells, clearly insufficient to explain the strong decrease in number (**Additional file 6**, C, lower panel). Altogether, these results indicate that PBMC culture media sustains live *M. bovis*, albeit at a very low replicating state, which is likely to reflect what happens in the context of a real infection.

### Ex vivo response to live Mycoplasmopsis bovis

Overall, the PBMC response to *M. bovis* differed to what observed in previous experiment employing TLR ligands, showing the importance to assess live pathogens rather than TLR ligands as surrogates of viral or bacterial infections. Indeed, this time the previously mentioned MHC-II down- regulation on monocytes or DCs did not occur, but the opposite (**Fig. 4A** and **4B**). Importantly, cells from comparators receiving heat-inactivated *M. bovis* as negative controls lacked clear evidence for specific activation/maturation, the normalized values for all investigated activation / maturation marker remaining stable at a value of 1 (grey zones, **Fig. 4A**, **4B** and **4C**). Overall, the most striking observation was a clear CD25 induction for all immune cell subsets, showing their tight cooperation in the response towards *M. bovis*.

**Figure 4.**
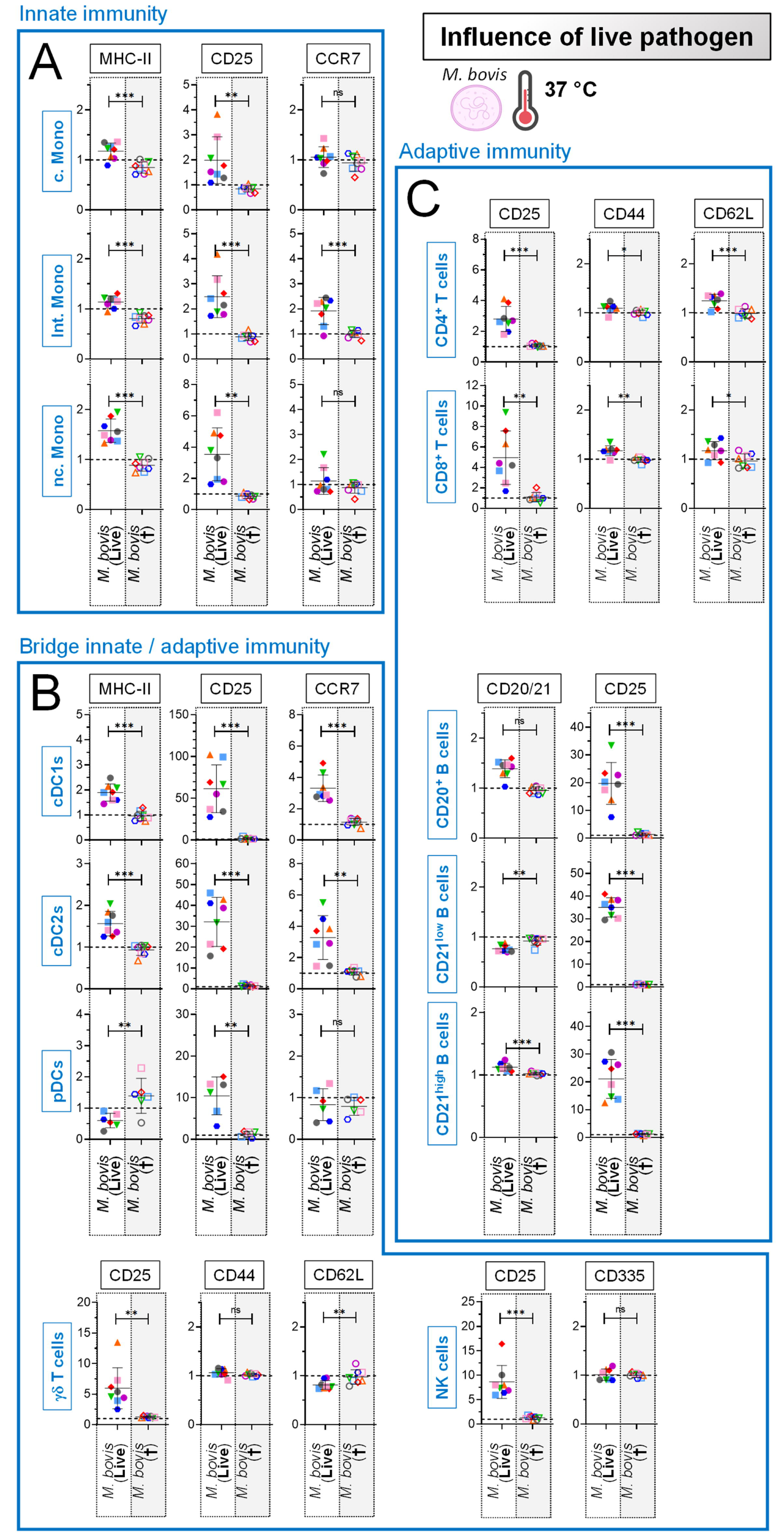
***Ex vivo* response to live *Mycoplasmopsis bovis.*** PBMCs from 8 individual animals were either for 18 h unstimulated (reference points for the assay) or stimulated with *M. bovis* (live or heat- inactivated). Stimulations were run at 37 °C. The fold changes analysis of activation/maturation markers was determined by FCM with FlowJo. Cells from the individual cows are represented by separate symbols; for stimulated samples, MFI values are normalized to that obtained with the reference point from the same animal. (**A**) Results obtained for the “Innate immunity” cell subsets. (**B**) Results obtained for the “Bridge Innate/adaptive immunity” cell subsets. (**C**) Results obtained for the “Adaptive immunity” cell subsets. The grey zones correspond to the cells from comparators receiving heat-inactivated *M. bovis*, as negative controls. Experimental conditions were compared using paired t tests. Stars indicate significance levels. *, p < 0.05; **, p < 0.01; ***, p < 0.001; ns: not significant.

### Natural killer cells are predominant IFN-γ secretors in response to *Mycoplasmopsis bovis*

Upon stimulation with live *M. bovis*, the most noticeable result at 37 °C consisted in increased levels of a set of pro-inflammatory cytokines, namely IL-1α, IL-1β, IL-6, MIP-1α, MIP-1β, and TNF-α, as well as pro-Th1 (IFN-γ) and pro-Th17 (IL-17). Of note, as observed previously for TLR ligand stimulation, the anti-inflammatory IL-10 paralleled this induction. Finally, and as for FCM results, stimulation by negative control (heat-inactivated *M. bovis*) had negligible effect on cytokine induction (**Fig. 5A**).

**Figure 5.**
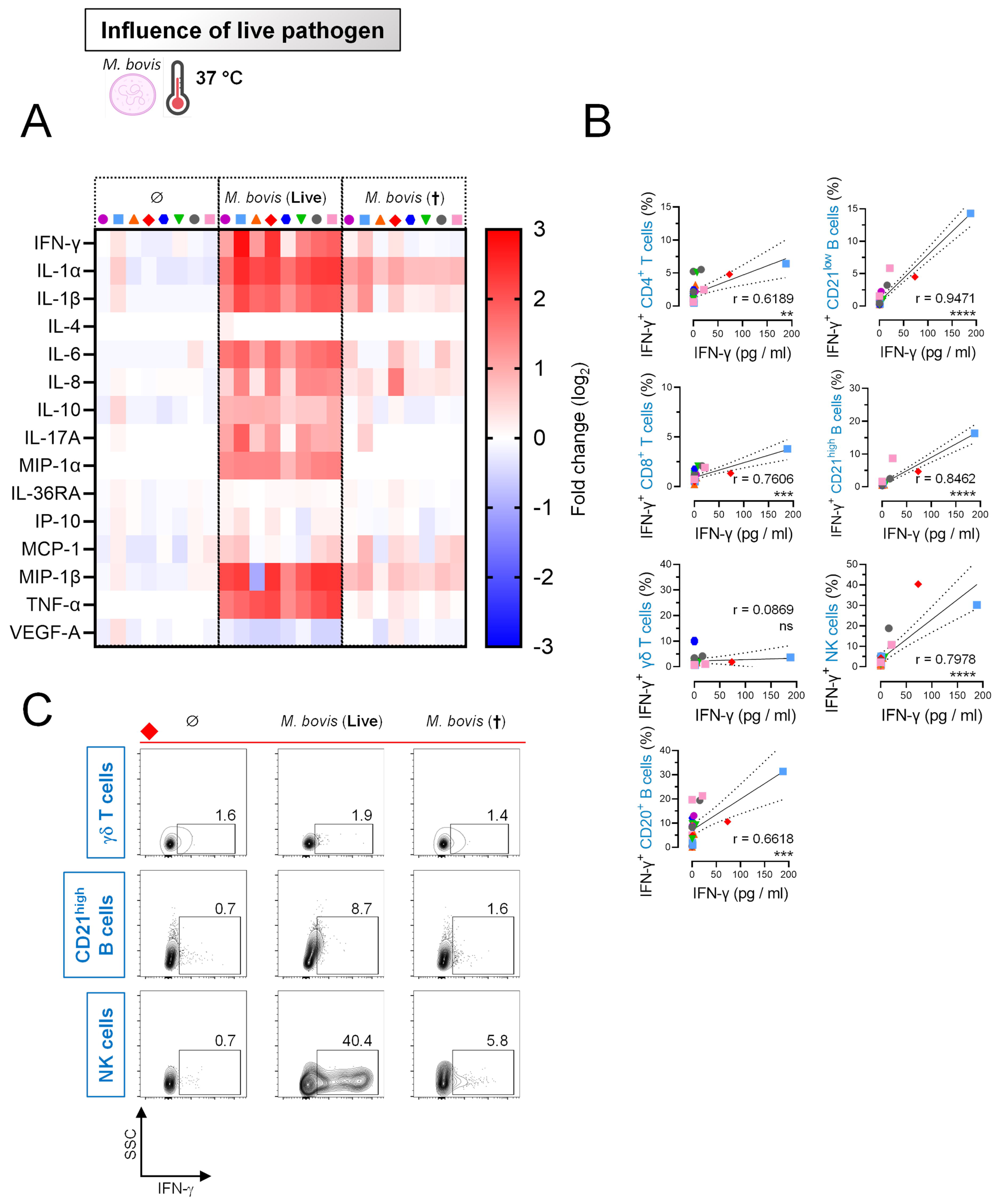
**Natural killer cells are predominant IFN-**γ **secretors in response to *Mycoplasmopsis bovis*.** PBMCs from 8 individual animals were either for 18 h unstimulated or stimulated with *M. bovis* (live or heat-inactivated). Stimulations were run at 37 °C. (**A**) Cytokine/chemokine secretion in supernatants of PBMC cultures, using Multiplex immunoassay. A single measurement was done per samples tested, and each symbol represents an individual cow. Heat map shows log_2_-fold changes in concentration of 15 Cytokines/chemokines. For a given cytokine/chemokine, normalization was as follow: [concentration for a given animal] / [average concentration of reference points for 8 animals]. **(B)** Correlation coefficient (r) obtained with percentage of IFN-γ^+^ cells for different immune cell subsets (measured by FCM) calculated as a function of IFN-γ concentration (pg/ml) in culture supernatant (measured by multiplex immunoassay). Significance was reached for all immune cell subsets, excepting γδ T cells. Each symbol represents an individual animal. Stars indicate significance levels. ***, p < 0.001; ****, p < 0.0001; ns: not significant. **(C)** FCM contour plots from a representative animal, gated on γδ T cells, CD21^low^ B cells or NK cells. Numbers indicate the percentage of IFN-γ^+^ cells for this specific animal. Notably, a clear activation (intracellular IFN-γ detection) of NK cells is found, with almost 40% of positive cells in optimal cell culture conditions.

Next, we wanted to ascertain that the concentrations of secreted cytokines (Multiplex immunoassay, (pg/ml)) were linked to the magnitude of cytokine production detected by FCM (intracellular cytokine staining (ICS), percentage of cytokine-producing cells). Moreover, unlike Multiplex immunoassay that was informative only on bulks of PBMCs, ICS offers the advantage to clarify which specific cell subsets produced individual cytokines. We opted for directly-labeled antibodies, since two-step ICS procedures implying secondary antibodies proved ineffective in our hands (data not shown). Four antibodies against IFN-γ, TNF-α, IL-4 and IL-17 were employed. We first ensured that our system could easy detect those cytokines following polyclonal stimulation of lymphocytes by concanavalin A or PMA/ionomycin (**Additional file 7**). Of note, bovine PBMCs were unable to produce detectable levels of IL-4 following *M. bovis* exposure (**Fig. 5A**), and IL-17 was hardly detectable by FCM, which is why this section focuses on IFN-γ and TNF-α. By combining results obtained with both procedures, we found a significant association between IFN-γ concentration and percentage of IFN-γ^+^ cells for most of investigated cell subsets, the best correlation being obtained for B and NK cells (**Fig. 5B**). This indicates that distinct subsets of bovine blood cells cooperate tightly with the magnitude of the humoral and cellular immune response against this *Mollicutes* pathogen. Remarkably, NK cells could elicit very high amount of IFN-γ, as illustrated in **Fig. 5C**.

When percentages of TNF-α^+^ cell subsets were plotted as a function of secreted TNF-α (Multiplex immunoassay), significant associations were also found; this time TNF-α production was mainly driven in coordinated manner by the monocyte, B and NK cell fractions (**Additional file 8**). Again, those results show that the combination of two complementing and independent techniques offers a powerful experimental setup for the best understanding of host-*M. bovis* interactions.

### Temperature impacts primary blood cell stimulation by *Mycoplasmopsis bovis*

Since the body temperature of many livestock species is generally at least 1-2 degrees above 37°C, we next investigated the influence of incubation temperature: 37 °C (standard cell culture conditions, “Hypothermia”) versus 38-38.5 °C (ruminant body temperature, “Body Temp”), postulating that the later would provide more relevant insights for understanding the interaction between host’s immune cell and *M. bovis*.

Strikingly, the rise of temperature proved to modulate the response to *M. bovis* of monocytes (MHC-II, CD25, and CCR7 downregulation, indicative of a reduced cellular activation) (**Fig. 6A**), γδ T cells (**Fig. 6B**), and T and B cells (reduced CD44 and CD25 expression, respectively) (**Fig. 6C**). Taken together, these results strongly suggest that the research work employing veterinary live pathogens should be run under temperature mimicking the hosts body temperature; otherwise, the commonly used incubation temperature of 37 °C, which is in a cow - strictly speaking - hypothermia, might lead to suboptimal – if not incorrect – immunological interpretations.

**Figure 6.**
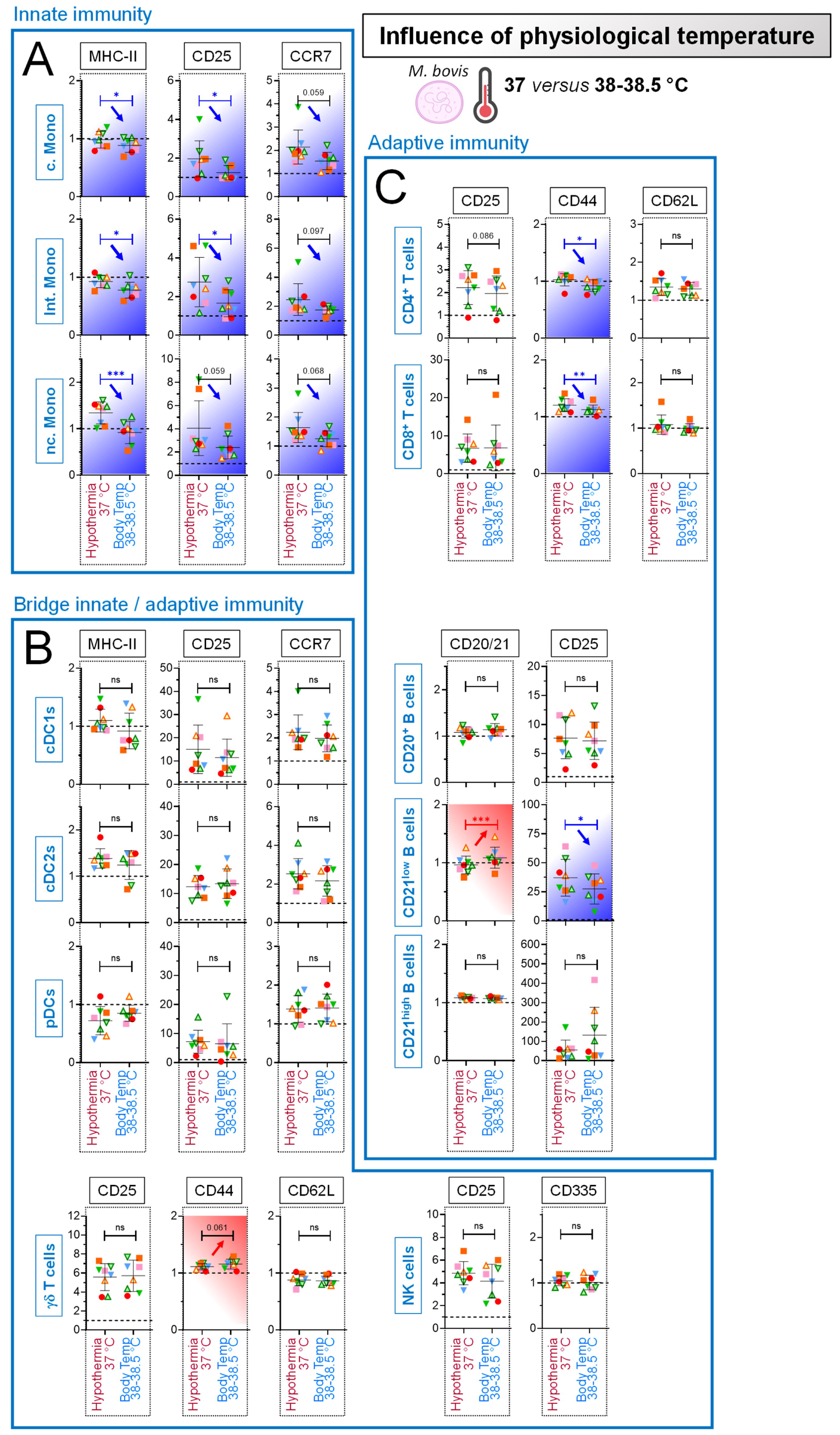
Influence of physiological temperature on primary blood cell stimulation by *Mycoplasmopsis bovis*. PBMCs from 8 individual animals were either for 18 h unstimulated (reference points for the assay) or stimulated with live *M. bovis*. Stimulations were run in parallel at 37 °C (standard cell culture condition, “Hypothermia”) or at 38-38.5 °C (physiological temperature of cow, “Body Temp”). The fold changes analysis of activation/maturation markers was determined by FCM with FlowJo. Cells from the individual cows are represented by separate symbols; for stimulated samples, MFI values are normalized to that obtained with the reference point from the same animal. (**A**) Results obtained for the “Innate immunity” cell subsets. (**B**) Results obtained for the “Bridge Innate/adaptive immunity” cell subsets. (**C**) Results obtained for the “Adaptive immunity” cell subsets. Red arrows indicate a further increase of activation/maturation marker expression induced by 38-38.5 °C compared to 37 °C; blue arrows indicate the opposite. Experimental conditions were compared using paired t tests. *, p < 0.05; **, p < 0.01; ***, p < 0.001; ns: not significant.

### Induction of cytokines by primary blood cells following exposure to *Mycoplasmopsis bovis* under physiological temperature

Again, it was considered important to corroborate the above results, which is why the PBMC cytokine secretion to *M. bovis* was evaluated. Firstly, when PBMCs were let unstimulated, the baseline levels of cytokines were poorly influenced by the rise of temperature to more physiological conditions (“Body Temp” (38-38.5 °C) versus “Hypothermia” (37 °C)).

Under bovine body temperature, *M. bovis* exposure also promoted a balanced cytokine response, consisting in a set of pro-inflammatory, pro-Th1, pro-Th17 and anti-inflammatory (IL-10) cytokines (**Fig. 7B**.) However, and in line with the notion that the overall activation/maturation state is modulated in more physiological temperatures, “Body Temp” gave an exacerbated and significant induction of IFN-γ, IL-6, IL-8 and IL17A, as shown in **Fig. 7B** (all animals depicted, pink to bright red patterns for all animals) and **Fig. 7C** (mean of eight animals). Altogether, this clearly showed that the use of physiological temperatures should be applied in the experimental design for the best characterization of pathogens- immune interactions.

**Figure 7.**
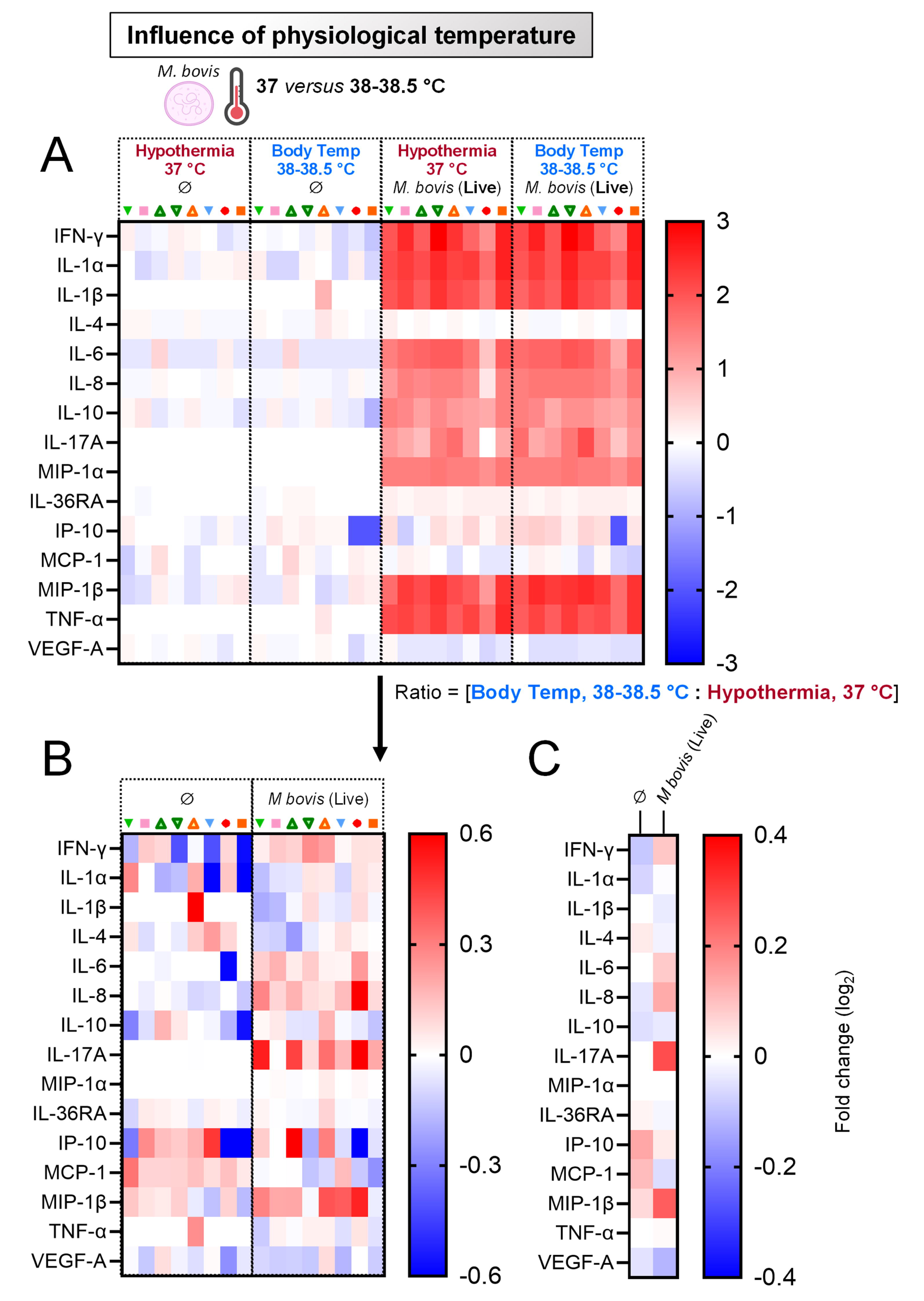
Induction of cytokines by primary blood cells following exposure to *Mycoplasmopsis bovis* under physiological temperature. PBMCs from 8 individual animals were either for 18 h unstimulated or stimulated with live *M. bovis*. Stimulations were run in parallel at 37 °C (standard cell culture condition, “Hypothermia”) or at 38-38.5 °C (“Body Temp”). (**A**) Cytokine/chemokine secretion in supernatants of PBMC cultures, using Multiplex immunoassay. A single measurement was done per samples tested, and each symbol represents an individual cow. Heat map shows log_2_-fold changes in concentration of 15 Cytokines/chemokines. For a given cytokine/chemokine, normalization was as follow: [concentration for a given animal] / [average concentration of reference points at 37 °C (“Hypothermia”, Ø) for 8 animals]. (**B**) As in (A), but displaying the ratios: [Body Temp : Hypothermia]. (**C**) As in (B), but showing average instead of individual values.

### Influence of fever on primary blood cell stimulation by *Mycoplasmopsis bovis*

The influence of fever was further analyzed, this time by raising the question of how stimulation at 38-38.5 °C (normal bovine body temperature, “No fever”), 39.5 °C (“Low fever”) and 41 °C (“high fever”) [29] would influence the magnitude of PBMC response to *M. bovis.* Indeed, fever is frequently a response to infection and immune responses occur commonly at fever temperatures, however this parameter remains poorly addressed. Importantly, results illustrated in **Additional file 9** clearly showed no impact of the rise from 38 °C to 41 °C on the percentage of living cells, regardless of whether the cells were unstimulated or stimulated with *M. bovis.* This was certainly not the case when the analysis was done for the activation/maturation markers. Indeed, “High fever” condition modulated up-regulation of MHC-II (positive effect on cDC1s), CCR7 (positive effect on cDC1s, negative effect monocytes) and CD25 (negative effect monocytes, DCs, B cells, T cells, γδ T cells and NK cells) (**Fig. 8A**, **8B** and **8C**). If a positive effect of “high fever” condition on the marker upregulation was seen in a few occasions (as it was the case for the cDC1 subset), the overall trend was a clear abrogation of the ability of immune subsets to maturate and activate in response to *M. bovis*, possibly preventing them to mount an efficient immune response.

**Figure 8.**
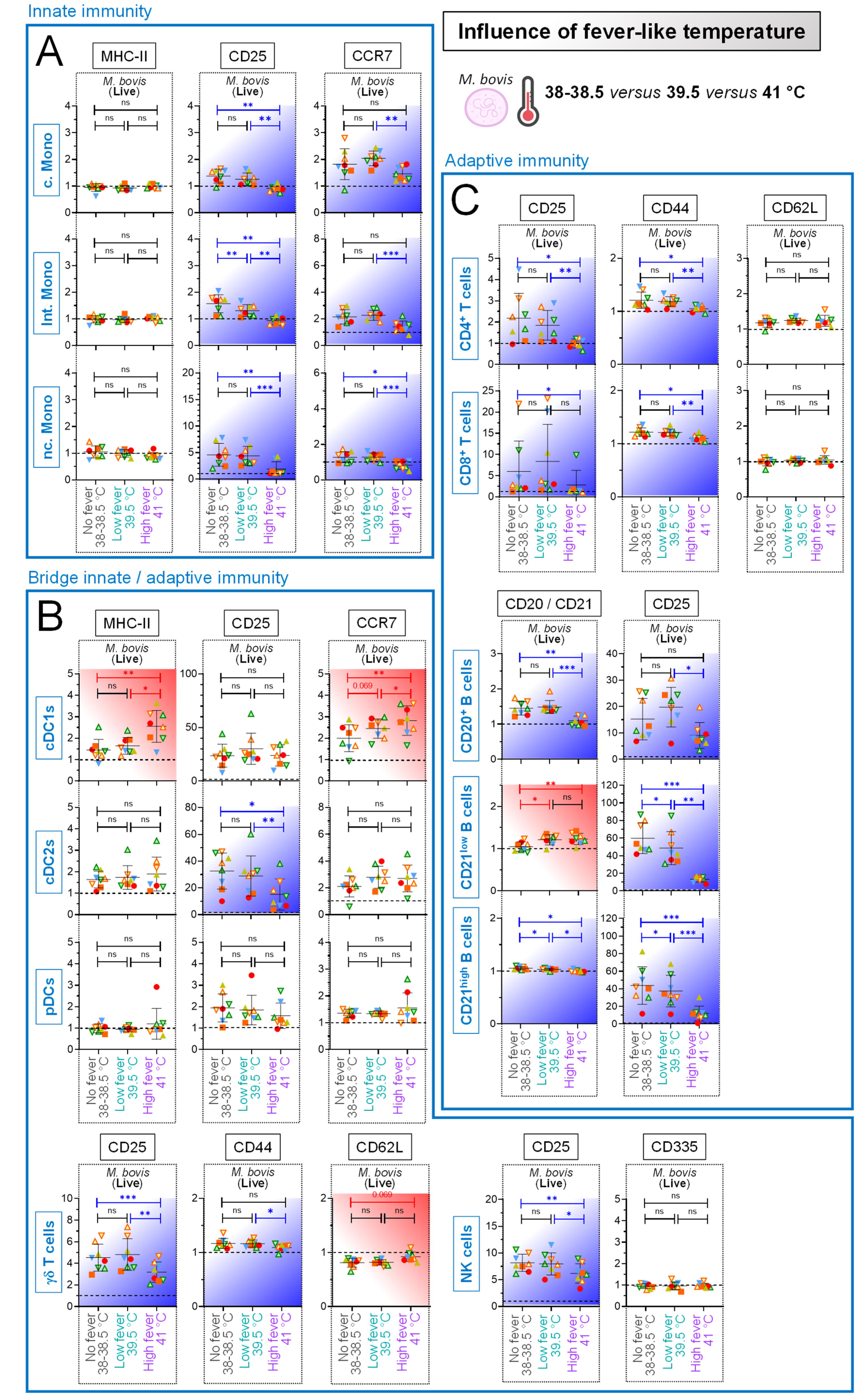
Influence of fever on primary blood cell stimulation by *Mycoplasmopsis bovis*. PBMCs from 8 individual animals were either for 18 h unstimulated (reference points for the assay), stimulated with live *M. bovis* at 38 °C (normal bovine body temperature, “No fever”), 39.5 °C (“Low fever”) or 41 °C (“High fever”). The fold changes analysis of activation / maturation markers was determined by FCM with FlowJo. Cells from the individual cows are represented by separate symbols; for stimulated samples, mean fluorescence intensity (MFI) values are normalized to that obtained with the reference point from the same animal. (**A**) Results obtained for the “Innate immunity” cell subsets. (**B**) Results obtained for the “Bridge Innate/adaptive immunity” cell subsets. (**C**) Results obtained for the “Adaptive immunity” cell subsets. Blue backgrounds indicate a significantly attenuated activation / maturation marker expression under high fever temperature, compared to low/no fever temperature; red backgrounds indicate the opposite. Experimental conditions were compared using one-way ANOVA followed by Tukey’s post hoc test. Stars indicate significance levels. *, p < 0.05; **, p < 0.01; ***, p < 0.001; ns: not significant.

### High fever reduces the secretion of pro-inflammatory cytokines by primary blood cells in response to *Mycoplasmopsis bovis*

In order to confirm the above results, we assessed PBMC cytokine response to *M. bovis* at those three different incubation temperatures. The rise from 38 °C to 41 °C had a minor influence on the overall cytokine secretion by unstimulated PBMCs. Again, regardless the incubation temperature, *M. bovis* exposure promoted a balanced cytokine response, comprising simultaneously pro-inflammatory (i.e. IL- 1α, IL-1β, IL-6, MIP-1α, MIP-1β, and TNF-α), pro-Th1 (IFN-γ) and anti-inflammatory (IL-10) cytokines. However, in line with the notion that the overall activation/maturation state was decreased, “High fever” condition led to an overall less robust response. This was witnessed by the reduced positive fold change (compared to “No fever” and “Low fever”, color patterns for most cytokines depicted in the “High fever” column turned pink rather than bright red) (**Fig. 9A**). Certainly, when the values obtained with “Low fever” were normalized to that obtained with normal bovine body temperature (“No fever”) (**Fig. 9B**), no influence was noted. In contrast, when the values obtained with “High fever” were normalized to that obtained with either normal bovine body temperature (“No fever”) (**Fig. 9C**) or “Low fever” (**Fig. 9D**), a very low intensity was calculated for all induced cytokines (blue color patterns), confirming that high fever condition strongly prevented PBMCs to secrete high levels of cytokine against *M. bovis*. This inhibition was particularly obvious for IFN-γ, a typical pro-Th1 cytokine. Collectively, these results highlighted the clear impact of temperature on PBMC capacity to be activated against pathogens such as *M. bovis*: optimal at normal body temperature (“No fever”) or low fever, strongly attenuated under conditions mimicking high fever temperatures.

**Figure 9.**
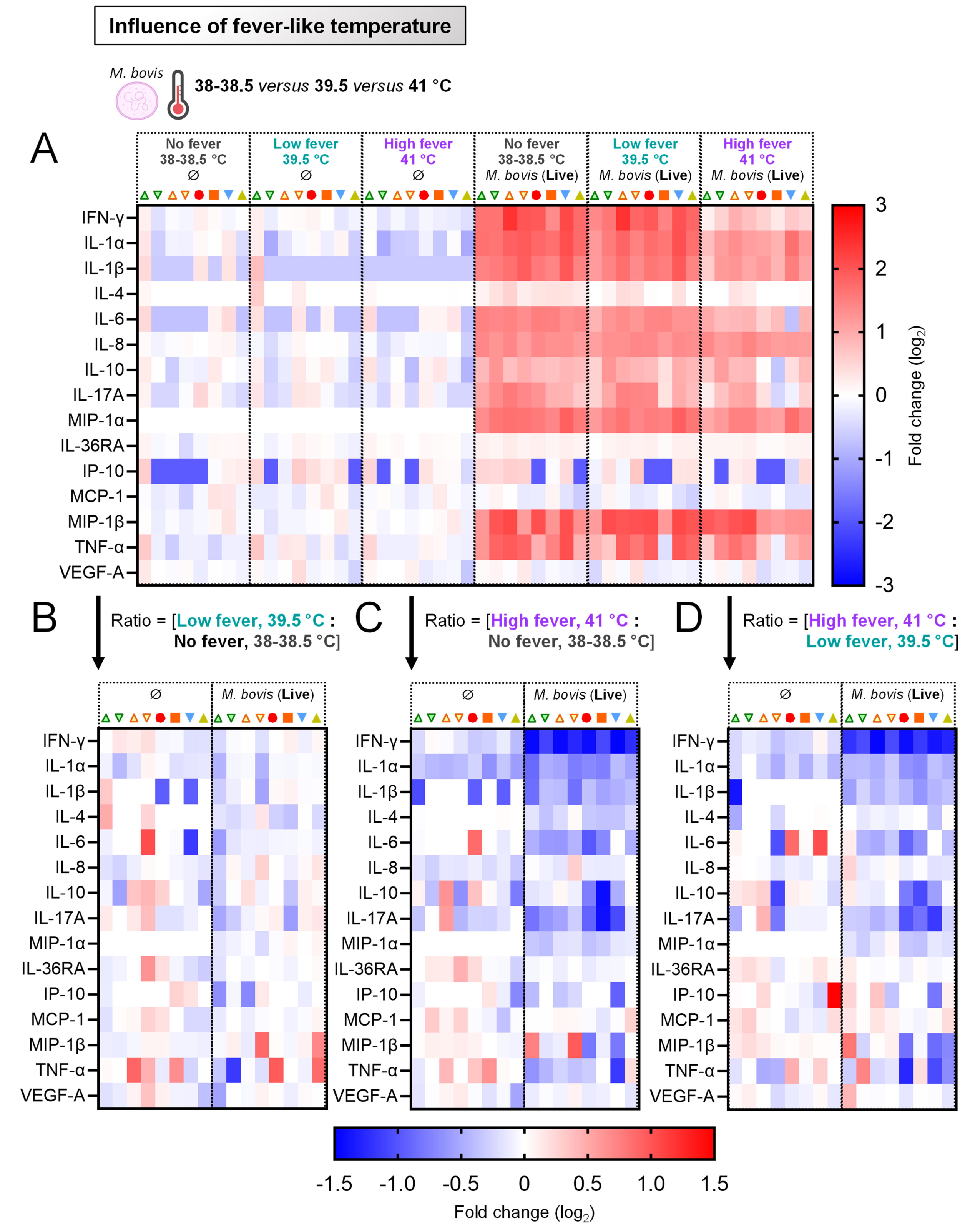
**High fever reduces the secretion of pro-inflammatory cytokines by primary blood cells in response to *Mycoplasmopsis bovis.*** PBMCs from 8 individual animals were either for 18 h unstimulated (reference points for the assay), stimulated with live *M. bovis* at 38 °C (normal bovine body temperature, “No fever”), 39.5 °C (“Low fever”) or 41 °C (“High fever”). (**A**) Cytokine/chemokine secretion in supernatants of PBMC cultures, using Multiplex immunoassay. A single measurement was done per samples tested, and each symbol represents an individual cow. Heat map shows log_2_-fold changes in concentration of 15 cytokines/chemokines. For a given cytokine/chemokine, normalization was as follow: [concentration for a given animal] / [average concentration of reference points at 38 °C (“No fever”, Ø) for 8 animals]. (**B**) As in (A), but displaying the ratios: [Low : No fever]. (**C**) As in (A), but displaying the ratios: [High : No fever]. (**D**) As in (A), but displaying the ratios: [High : Low fever].

## DISCUSSION

To properly assess the concerted actions of immune cells against pathogens, it is essential to use an experimental set-up that encompasses the majority of immune cell subsets. Our *ex vivo* platform employed freshly isolated PBMCs and therefore major subsets of immune cells. The use of PBMCs is in line with the 3R principles, associated with easy sampling and the advantage to be collectable from a number of different animals on a frequent basis associated with only minimal distress. Clearly, *in vivo* studies are most informative, but challenging in terms of standardization, costs associated with staff as well as containment and most importantly they negatively impact the wellbeing of animals. Studies conducted with PBMCs are a good surrogate model for understanding the initial phase of host-pathogen interactions involving cells of the innate arm of immunity [30] – particularly for pathogens that go through a bacteremia phase as it is the case of *M. bovis*. Interaction of *M. bovis* with PBMCs were already reported, but with inconsistent findings regarding host cells’ viability, maturation and activation [31–35]. Consequently, when initiating this study, an unequivocal description of *M. bovis* interaction with PBMCs was still missing to reconcile those discrepancies. Finally, we speculated that the invasion of circulating immune cells by *M. bovis* could contribute to pathogenesis by helping pathogen dissemination from the respiratory tract to other sites such as joints or the udder.

However, the use of PBMCs, like every other model system, has limitations. Our system does not constitute a good model to evaluate long-term outcomes of infections *per se*. Effectively, the development of an immunological memory implies cell migration to draining LNs, and this could not be evaluated in the present study. In line with this statement, our results obtained with CD44 and CD62L markers (allowing the clear distinction between naïve and memory T cells [36]) were less informative and lowly modulated in comparison to CD25. Nonetheless, their inclusion in the “T cells” panel was considered crucial, having in mind that this experimental platform might be also applied to assays employing blood from animals which have been vaccinated or challenged. Of note, when this study was initiated and conducted, CD45RO antibody commonly used to identify memory T cells [37] was discontinued, with the consequence that we had to use CD44 instead, which has known limitations.

Taking advantage of the unprecedented delineation of bovine immune components offered by our platform, we intended at first to revisit part of the current knowledge, as well as providing new insights in interactions of bovine immune cells with *M. bovis* (**Figs. 4**-**5**). Concerning the monocyte subset, we saw only a moderate modulation of cell surface markers, with no obvious up-regulation of MHC-II (related to antigen presentation) and CCR7 (related to cell migration). However, a clear TNF-α induction was observed, showing that monocytes act locally to promote a pro-inflammatory environment. Moreover, we provided the direct evidence that they are the main source of TNF-α within PBMCs, when previous reports looked at TNF-α production only on a bulk of PBMCs [33] or purified monocytes [2]. To date, the function of the DC subset upon *M. bovis* infections has not been directly investigated yet. We quantified a robust MHC-II, CD25 and CCR7 upregulation on both cDC1 and cDC2 subsets, which strongly suggest a typical behavior of those APCs: emigrate to LNs whereby they can process *M. bovis* antigens. Potential role of NK cells in the host defense against *M. bovis* – and more generally *Mollicutes* – remains under investigation, although sparse data suggest their involvement, possibly linked to the severity of infection [31, 38]. Remarkably, our data showed a spectacular IFN-γ production by NK cells, making from now unequivocal the tight interaction of this cell subset with *M. bovis*. Lastly, we found that γδ T cells contributed to the overall response (as witnessed by the CD25 upregulation), but we could not attribute them a particular role despite their overrepresentation in cattle, in line with a previous study questioning their specific function [39]. Of note, γδ T cells were identified by the expression of WC1 which does not encompass the whole subset circulating in peripheral blood, and the use of an anti-δ chain specific antibody might have resulted in a slightly different picture [40, 41]. Consequently, there is still some debate whether γδ T cells ultimately participate in bacterial clearance or immunopathology.

As mentioned above, the peripheral blood is not the main compartment for initiating adaptive immunity, while it contains clearly CD4^+^ and CD8^+^ T cell subsets. Effectively, exposure to *M. bovis* led to a little modulation of CD25 expression and had no effect on CD44 and CD62L markers. Nonetheless, activation signs were clearly observed, witnessed by the apparition of IFN-γ-producing T cells, in line with a previous study [31]. This was most likely a bystander effect – activation of non-specific T cells without cognate antigens –, as this phenomenon is known to happen during bacterial infection [42, 43]. Another remarkable observation was the elevated amount of the anti-inflammatory IL-10, cytokine notably driven by Treg cells, whose production is expected to occur at a later step of the sequential immune response. This could be an immune evasion strategy of *M. bovis* via the secretion of a 26-amino-acid peptide; in the end this would promote the shift to a Th2 cytokine profile, with the subsequent diminishing of opsonization capacity [2, 44]. Finally, no pan B cell markers are known in cattle, and this is only very recently, that putative B cell populations were proposed [45]. This is why we made a distinction between CD20^+^, CD21^low^ and CD21^high^ cells – the last two subsets having greater chance to constitute the memory B cell pool [46]. Again, activation signs were seen, the most noticeable being IFN-γ and TNF-α secretion by both CD21^low^ and CD21^high^ B cell populations. Since memory but not naïve B cells can produce TNF-α [47], it really indicated that our gating strategy considering the inclusion of CD21^low^/^high^ was adequate to phenotype memory B cells. In conclusion, we were not only able to recapitulate the features of *M. bovis* immunity in great details, but we also provided novel insights into the tight cooperation of the host’s contributing factors, including the so far overlooked NK cell subset. Collectively, these findings will be of great help to better understand the orchestrated immune responses against *Mollicutes*, which in turn, would benefit to the development of rationally designed vaccines.

The shift from 37 °C (“Hypothermia”) to a temperature more related to ruminant body temperature had immunological consequences by affecting the magnitude of the PBMC response to *M. bovis* (**Figs. 6**- **7**). These important findings should encourage the research community to perform immunological assays of ruminants at physiological body temperature, as it is done for years in the porcine field with experiments run at 39 °C [48–50]. We then sought to assess whether a temperature mimicking high fever would exacerbate this PBMC response (**Figs. 7**-**8**). Indeed, fever is a hallmark of infection and inflammatory disease [51], and typical *M. bovis* infection symptoms include fever for at least several days. The fact that fever was conserved through evolution strongly argues that it confers a benefit in clearing infections; but how this acts exactly to clear invading pathogens is yet still unclear. In addition, in cases of extreme inflammation, lowering rather than raising body temperature was reported to be protective [52]. Similarly, high fever is associated with a greater incidence of serious bacterial diseases such as pneumonia or sepsis [53]. In this study, the shift of temperature to mimic high fever conditions had effectively drastic effects, giving rise to the more intriguing results. Indeed, far from inducing an overall increased PBMC response, this time we obtained a significant reduction of maturation/activation parameters upon interaction with *M. bovis*. This, regardless of cell subsets considered, excepting cDC1s. This corroborates studies available for decades, whereby elevated temperatures substantially enhanced DC function, including MHC and co-stimulatory molecules, as well as migration capacity [54–57]. In our case, whether this increased DC function can compensate the attenuation of all other subsets to mount an efficient adaptive immune response remains to be assessed, although quite unlikely.

Several factors can explain this high fever mediated immunomodulation. One might argue that PBMCs are perishable living cells, some of them beginning to die immediately after their isolation from whole blood. Thus, our cultured PBMCs might be particularly sensitive to high fever-like temperatures – more than their counterparts in physiological conditions –, as previously reported [58, 59]. Nevertheless, our results clearly showed that viable cell frequencies were unaffected at elevated temperatures, contradicting this hypothesis. It would be also possible that 41 °C affects the pathogen, explaining why we obtained intermediate readouts between what measured for heat-inactivated (no PBMC response) or live *M. bovis* stimulation (strong PBMC response). There is little chance that it was the case, since *M. bovis* was shown to be protected from temperature changes (up to 50 °C) for an extended period of time via the production of a biofilm [60]. Another explanation would be that fever temperatures are likely to promote a Th2 switch in responding CD4 T cells, possibly modulated further by IL-12 [61]. Due to technical limitations (investigation limited to blood compartment from naïve animals; restricted cytokine detection tools in cattle), the eventual Th2 skewing hypothesis remained speculative in our model. Effectively, despite the strong reduction of IFN-γ secretion under high fever condition, we failed to detect instead any consistent IL-4. Moreover, we were lacking the possibility to detect any IL-12 and IL-13 (another pro-Th2 cytokine), those being absent from the bovine multiplex immunoassay. Altogether, our results suggest that high fever is associated with worse outcome in animal infected with *M. bovis,* which has been shown experimentally for other *Mollicutes* infecting ruminants [62, 63].

In conclusion, our experimental platform allowed an unprecedented phenotypic delineation of bovine immune responses in the context of *M. bovis*. This platform has a great prospect to decipher concerted actions of immune cell subsets. Altogether, these results constitute a significant step in the better understanding of *M. bovis* immunity, with new insights related to the influence of low and high fever to combat infection. As a next step, we intend to perform coinfection experiments to check the synergism of *M. bovis* and other infectious pathogens (including vector-borne viruses such as Bluetongue and Schmallenberg viruses), since *M. bovis* can exacerbate respiratory disease to other pathogens. In future studies we will also use the platform to decipher host-pathogen interactions of different cattle breeds.

## DECLARATIONS

### Ethics approval

The collection of bovine blood was performed in compliance with the Swiss animal protection law (TSchG SR 455; TSchV SR 455.1; TVV SR 455.163) under the cantonal license BE55/2022. The experiments were reviewed by the cantonal committee on animal experiments of the canton of Bern and the Fribourg Commission, Switzerland, and approved by the cantonal veterinary authority (Amt für Landwirtschaft und Natur LANAT, Veterinärdienst VeD, Bern, Switzerland).

### Consent to participate

Not applicable

### Availability of data and materials

Whole genome sequences of 21 cows are available at NCBI Sequence Read Archive (PRJNAxxxxxx; pending).

### Competing interests

The authors declare no competing financial interests.

### Funding

The work was funded by the European Union’s Horizon 2020 research and innovation programme PREPARE4VBD (grant No 101000365) and received also support from the Swiss National Science Foundation (grant number 310030_201152, www.snf.ch).

### Authors’ contributions

All authors contributed important elements to the work presented in this paper.

T.D. designed and performed the experiments, assembled the data, prepared the figures and drafted the manuscript.

T.Y. performed the experiments and edited the manuscript.

D.L. performed the genomic analysis and edited the manuscript.

F.L. performed M. bovis titer determination and colony micrograph, provided helpful discussion and edited the manuscript.

L.E. and R.S. organized the animal blood collection and edited the manuscript.

J.J. designed the overall concept and edited the manuscript.

## Supporting information

Characteristics of Animals

Antibodies against M. bovis were not detected in the serum of animals enrolled in the study

Heatmap of pairwise kinship estimates from genomic data

First and second principal components (eigenvector 1 and 2) estimated from genomic data by principal component analysis (PCA)

FCM gating strategy for immune cell identification

Mycoplasmopsis bovis survival in primary blood cell culture medium

Induction of cytokines by primary blood cells following exposure to mitogens

TNF-alpha production in response to Mycoplasmopsis bovis is a concerted action of monocytes, B cells and NK cells

Impact of the rise of temperature on viable cell frequencies

## Acknowledgements

We thank Stephanie Talker and Alizée Huber for precious advice with the Cytek Aurora; Sergi Torres-Puig for helpful discussions; Lutz Merbold and Yvo Aeby for initiating the collaboration with Agroscope animal facilities; Nicolas Ruggli, Roman Troxler, Jan Salchli and Hans-Peter Lüthi for blood collection.

## SUPPLEMENTARY FIGURES

Additional file 1. Characteristics of Animals

**Additional file 2. Antibodies against M. bovis were not detected in the serum of animals enrolled in the study.**

**Additional file 3. Heatmap of pairwise kinship estimates from genomic data.** The dendrogram depicts clusters identified by the complete linkage method of the hclust function in R. Colors indicate relatedness according to KING cutoff values as duplicates/monozygotic twins (> 0.354; red), 1^st^-degree (0.177 – 0.354; orange), 2^nd^-degree (0.0884 – 0.177; yellow), or 3^rd^-degree relatives (0.0442 – 0.0884; light blue), and more distantly related/unrelated individuals (< 0.0442; blue).

Additional file 4. First and second principal components (eigenvector 1 and 2) estimated from genomic data by principal component analysis (PCA). **Symbols represent individual cows.**

**Additional file 5. FCM gating strategy for immune cell identification.** (**A**) *Example of gating strategy for multiparameter FCM analysis of bovine DCs*. Gating strategy following multiparameter FCM staining uses Abs against MHC-II, CD172a, CD16, CD14, CD13, CD4, CD11c, CD25 and CCR7. A primary gate (P1) was set on FSC-A versus SSC-A. Then, the dead cell population (positive for Live/Dead) was excluded, followed by a FSC-H/FSC-A contour plot (to exclude doublets from the analysis). Among these cells, we defined the population positive for MHC-II and CD172a (low and intermediate only). Then, we defined among this population the cells negative for both CD14 and CD16 markers. Then, we defined among this population either the cells positive for CD13 and negative for CD4 markers (pre- cDC1s), either cells negative for CD13 and positive for CD4 markers (pre-pDCs), either cells negative for both (pre-cDC2s). Next, we considered CD11c marker to gate cDC1s and cDC2s (respectively pre-cDC1s and pre-cDC2s positive CD11c), as well as pDCs (pre-pDCs negative for CD11c). Maturation was evaluated based on the MFI of surface expression for MHC-II, CD25 and CCR7. (**B**) *Example of gating strategy for multiparameter FCM analysis of bovine monocytes*. Gating strategy following multiparameter FCM staining uses Abs against MHC-II, CD172a, CD16, CD14, CD13, CD4, CD11c, CD25 and CCR7. A primary gate (P1) was set on FSC-A versus SSC-A. Then, the dead cell population (positive for Live/Dead) was excluded, followed by a FSC-H/FSC-A contour plot (to exclude doublets from the analysis). Next, we defined among this population the cells highly positive for CD172a marker (CD172a^high^ cells), positive for CD13, and then negative for CD4. From this, classical monocytes (c.Mono) were gated as CD14^+^CD16^-^, intermediate monocytes (int.Mono) as CD14^+^CD16^+^, and nonclassical monocytes (nc.Mono) as CD14^-^CD16^+^. As for the DC substets, maturation was evaluated based on MFI of surface expression for MHC-II, CD25 and CCR7. White peak correspond to the signal obtained for MFO. (**C**) *Example of gating strategy for multiparameter FCM analysis to identify bovine T cells.* Gating strategy uses Abs against CD3, CD4, CD8, CD25, WC1, CD44, CD62L, IL-4, IFN-γ and IL-17. A primary gate (P1) was set on FSC-A versus SSC-A encompassing both small and large cells. Then, the dead cell population (positive for Live/Dead) was excluded, followed by a FSC-H/FSC-A contour plot (to exclude doublets from the analysis). Then, we considered cells positive for CD3, and defined among this population the cells negative for both CD4 and CD8 markers. Among these cells, γδ T cells were defined as the WC1^+^ cells, and the maturation was evaluated based on CD25 expression, whereas activation was evaluated on the basis of IFN-γ and/or IL-17. Alternatively, we defined among the CD3^+^ population either the cells positive for CD4 and negative for CD8 markers (CD4^+^ T cells), either cells positive for CD8 and negative for CD4 markers (CD8^+^ T cells). The maturation was evaluated based on CD25 expression, as well as percentage of central (CD62L^+^CD44^+^) or effector (CD62L^-^CD44^+^) memory cells. T cell subset activation was evaluated based on IL-4, IL-17 and IFN-γ production. (**D**) *Example of gating strategy for multiparameter FCM analysis of bovine NK cells*. Gating strategy following multiparameter FCM staining uses Abs against CD3, CD20, CD21, CD335 and CD25. A primary gate (P1) was set on FSC-A versus SSC-A. Then, the dead cell population (positive for Live/Dead) was excluded, followed by a FSC-H/FSC-A contour plot (to exclude doublets from the analysis). Next, we defined among this population the cells negative for CD3 marker. Then, we defined among this population the cells negative for both CD20 and CD21 markers. From this, NK cells were gated as CD335^+^ cells, and the maturation was evaluated based on CD25 expression, whereas activation was evaluated on the basis of IFN-γ, TNF-α and IL-17 production.

**Additional file 6: *Mycoplasmopsis bovis* survival in primary blood cell culture medium. (A)** Colonies with typical *M. bovis* morphology (Donetta PG45 strain, used in the present study). The size bar displays 1,000 μm. **(B)** CFU counts taken at different time points. *M. bovis* was grown in MEX or DMEM/10%, either at 37 °C (Hypothermia) or 39.5 °C (Low fever). **(C)** Similar as (B), but with FCM as readout. To determine the percentage of live *M. bovis*, *Bac*Light^TM^ RedoxSensor^TM^ Green Vitality Kit was employed.

**Additional file 7: Induction of cytokines by primary blood cells following exposure to mitogens.** (**A**) Cytokine / chemokine secretion in supernatants of PBMC cultures, using Multiplex immunoassay. Stimulations were run at 37 °C. A single measurement was done per samples tested, and each symbol represents an individual cow. Heat map shows log_2_-fold changes in concentration of 15 cytokines/chemokines. For a given cytokine / chemokine, normalization was as follow: [concentration for a given animal] / [average concentration of reference points (Ø) for 8 animals]. (**B**) As in (A), but displaying the ratios: [PMA/Iono : ConA]. (**C**) Here are shown IL-17 producing cells among the γδ T cell fraction (FCM contour plots) for 4 animals. Numbers indicate the percentage of IL-17^+^ cells.

**Additional file 8. TNF-**α **production in response to *Mycoplasmopsis bovis* is a concerted action of monocytes, B cells and NK cells. (A)** Correlation coefficient (r) obtained with percentage of TNF-α^+^ cells for different immune cell subsets (measured by FCM) calculated as a function of TNF-α concentration (pg/ml) in culture supernatant (measured by multiplex immunoassay). Each symbol represents an individual animal. Stars indicate significance levels. ***, p < 0.001; ****, p < 0.0001. **(B)** FCM contour plots from a representative animal, gated on TNF-α-producing cells (classical monocytes and NK cells, where a correlation was found) and non-producing cells (CD20^+^ B cells, where no correlation was found). Numbers indicate the percentage of TNF-α^+^ cells for this specific animal.

**Additional file 9. Impact of the rise of temperature on viable cell frequencies.** A primary gate (P1) was set on FSC-A versus SSC-A, then, the percentage of living PBMCs (negative for Live/Dead marker) was quantified by FCM. Cells were let either unstimulated (Ø), either stimulated with *M. bovis* at 3 different incubation temperatures: 38 °C (“No fever”), 39.5 °C (“Low fever”) and 41 °C (“High fever”). Each symbol represents an individual animal and is the average of three independent measurements (“Antigen presenting cells”, “T cells”, and “B cells, NK cells” panels). ns: not significant.

## Notes

### Competing Interest Statement

The authors have declared no competing interest.

