## Supplementary material for "Temperature effects on the magnitude of the bovine *ex vivo* immune response towards *Mycoplasmopsis bovis*": Characteristics of Animals

| Animal | Location | Breed | ID | Age (years) | Sex |
|---|---|---|---|---|---|
| 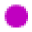 | Agroscope | *Bos taurus* | CH120150276437 | 2 | Female |
| 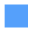 | Agroscope | *Bos taurus* | CH120150276420 | 2 | Female |
| 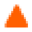 | Agroscope | *Bos taurus* | CH120150276581 | 1 | Female |
| 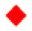 | Agroscope | *Bos taurus* | CH120150276451 | 2 | Female |
| 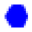 | Agroscope | *Bos taurus* | CH120150276444 | 2 | Female |
| 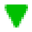 | Agroscope | *Bos taurus* | CH120150276499 | 1 | Female |
| 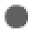 | Agroscope | *Bos taurus* | CH120150276482 | 1 | Female |
| 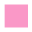 | Agroscope | *Bos taurus* | CH120150276475 | 1 | Female |
| 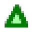 | Agroscope | *Bos taurus* | CH120150276550 | 1 | Female |
| 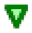 | Agroscope | *Bos taurus* | CH120150276567 | 1 | Female |
| 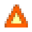 | Agroscope | *Bos taurus* | CH120150276512 | 1 | Female |
| 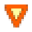 | Agroscope | *Bos taurus* | CH120158564574 | 1 | Female |
| 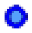 | Agroscope | *Bos taurus* | CH120150276574 | 1 | Female |
| 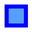 | Agroscope | *Bos taurus* | CH120150276635 | 1 | Female |
| 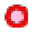 | Agroscope | *Bos taurus* | CH120150276628 | 1 | Female |
| 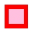 | Agroscope | *Bos taurus* | CH120158564628 | 1 | Female |
| 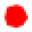 | Agroscope | *Bos taurus* | CH120150276505 | 2 | Female |
| 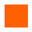 | Agroscope | *Bos taurus* | CH120150276611 | 2 | Female |
| 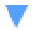 | Agroscope | *Bos taurus* | CH120150276543 | 2 | Female |
| 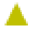 | Agroscope | *Bos taurus* | CH120150276529 | 2 | Female |
| 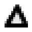 | Agroscope | *Bos taurus* | CH120150276581 | 1 | Female |
| 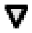 | Agroscope | *Bos taurus* | CH120150276468 | 2 | Female |
