## Supplementary material for "Temperature effects on the magnitude of the bovine *ex vivo* immune response towards *Mycoplasmopsis bovis*": Antibodies against M. bovis were not detected in the serum of animals enrolled in the study

A

**Raw data**

| | 1 | 2 | 3 | 4 | 5 | 6 | 7 | 8 |
|---|---|---|---|---|---|---|---|---|
| **A** | 0.066 | 0.065 | 0.104 | 0.057 | 0.082 | 0.057 | 0.07 | 0.071 |
| **B** | 0.039 | 0.092 | 0.09 | 0.059 | 0.086 | 0.06 | 0.073 | 0.073 |
| **C** | 1.276 | 0.066 | 0.079 | 0.085 | 0.063 | 0.29 | 0.092 | 0.071 |
| **D** | 1.154 | 0.063 | 0.1198 | 0.092 | 0.049 | 0.078 | 0.099 | 0.078 |
| **E** | 0.082 | 0.111 | 0.158 | 0.082 | 0.228 | 0.084 | 0.114 | 0.085 |
| **F** | 0.057 | 0.085 | 0.12 | 0.108 | 0.073 | 0.079 | 0.079 | 0.069 |
| **G** | 0.047 | 0.066 | 0.049 | 0.055 | 0.059 | 0.091 | 0.093 | |
| **H** | 0.055 | 0.083 | 0.046 | 0.043 | 0.058 | 0.076 | 0.07 | |

B

**Analysed data**

| | 1 | 2 | 3 | 4 | 5 | 6 | 7 | 8 |
|---|---|---|---|---|---|---|---|---|
| **A** | | | | | | | | |
| **B** | **0.0** | 2.2 | 3.8 | 0.5 | 2.7 | 0.5 | 1.6 | 1.7 |
| **C** | | | | | | | | |
| **D** | **100.0** | 1.0 | 4.0 | 3.1 | 0.3 | 11.3 | 3.7 | 1.9 |
| **E** | | | | | | | | |
| **F** | 1.5 | 3.9 | 7.4 | 3.7 | 8.4 | 2.5 | 3.8 | 2.1 |
| **G** | | | | | | | | |
| **H** | -0.1 | 1.9 | -0.4 | -0.3 | 0.5 | 2.7 | 2.5 | |
