## Supplementary figures and images for "Temperature effects on the magnitude of the bovine *ex vivo* immune response towards *Mycoplasmopsis bovis*"

### FCM gating strategy for immune cell identification

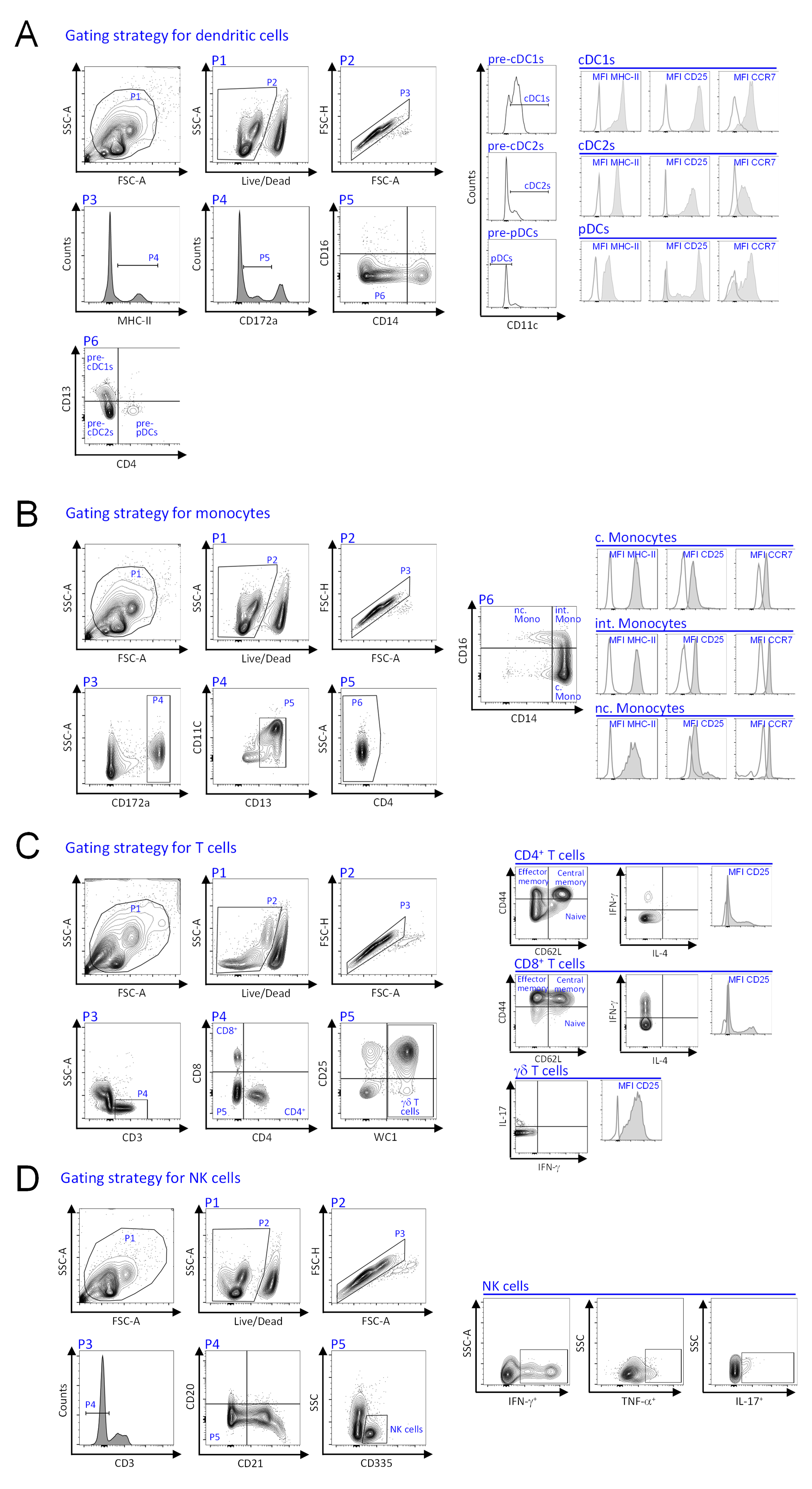

### First and second principal components (eigenvector 1 and 2) estimated from genomic data by principal component analysis (PCA)

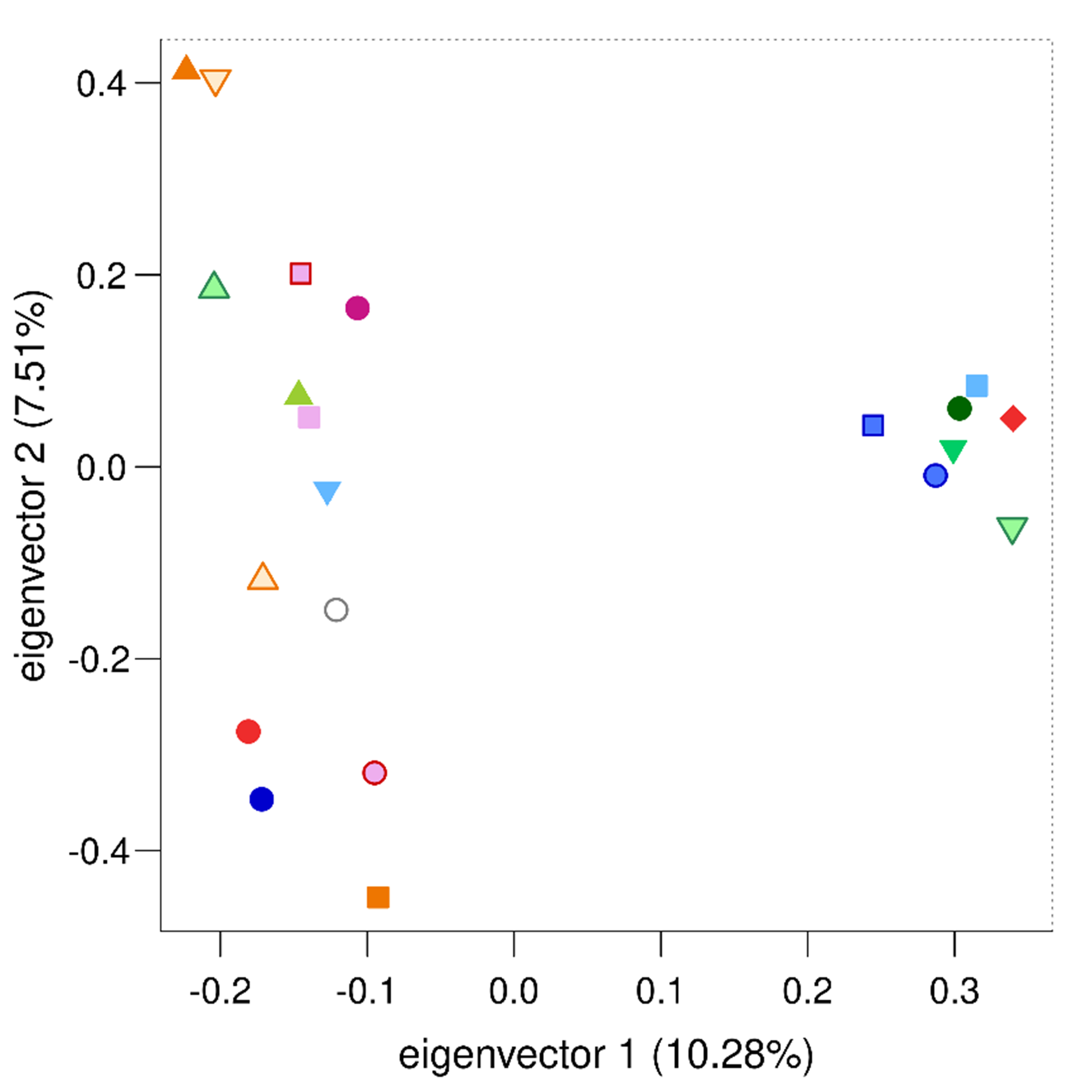

### Heatmap of pairwise kinship estimates from genomic data

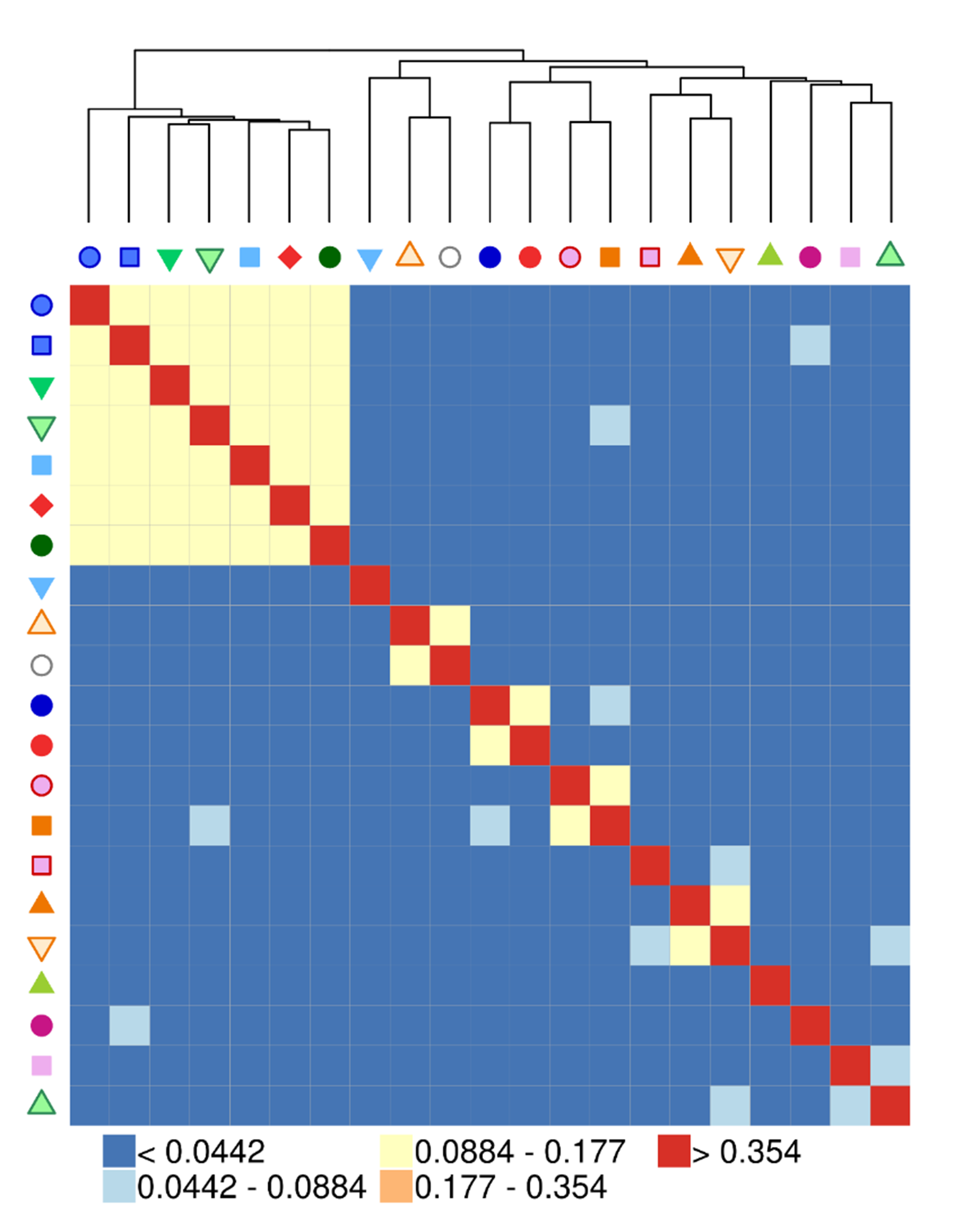

### Impact of the rise of temperature on viable cell frequencies

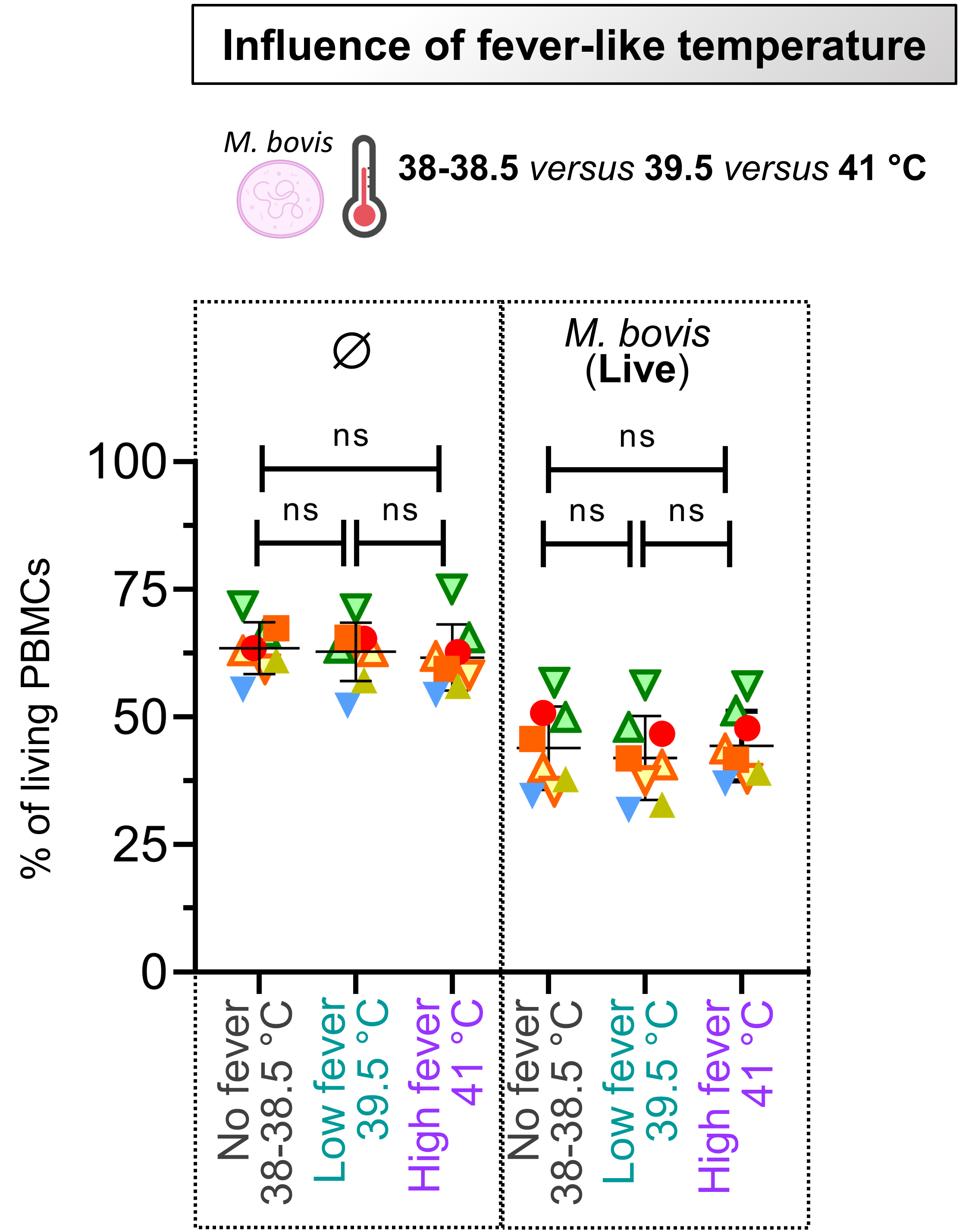

### Induction of cytokines by primary blood cells following exposure to mitogens

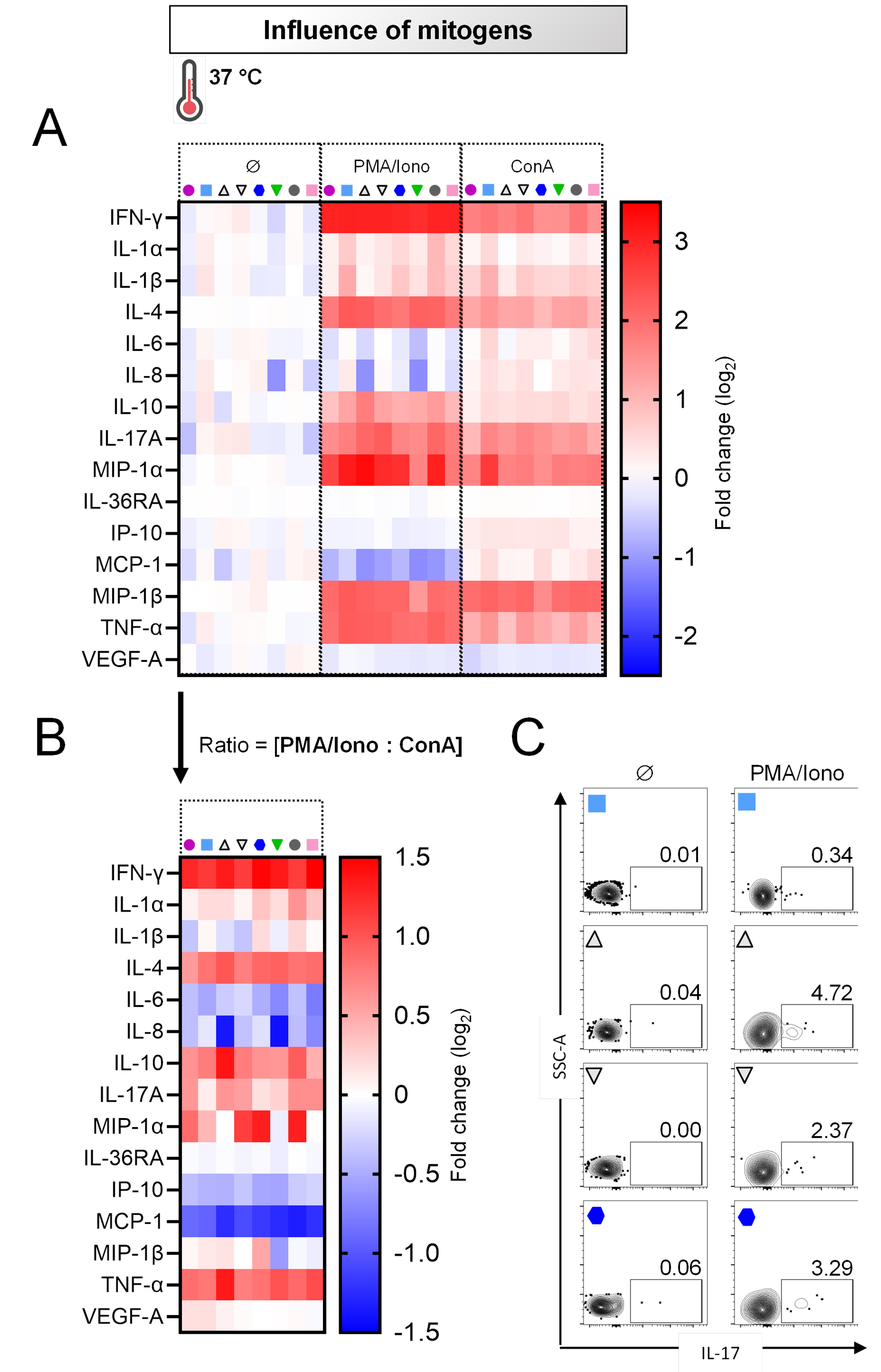

### Mycoplasmopsis bovis survival in primary blood cell culture medium

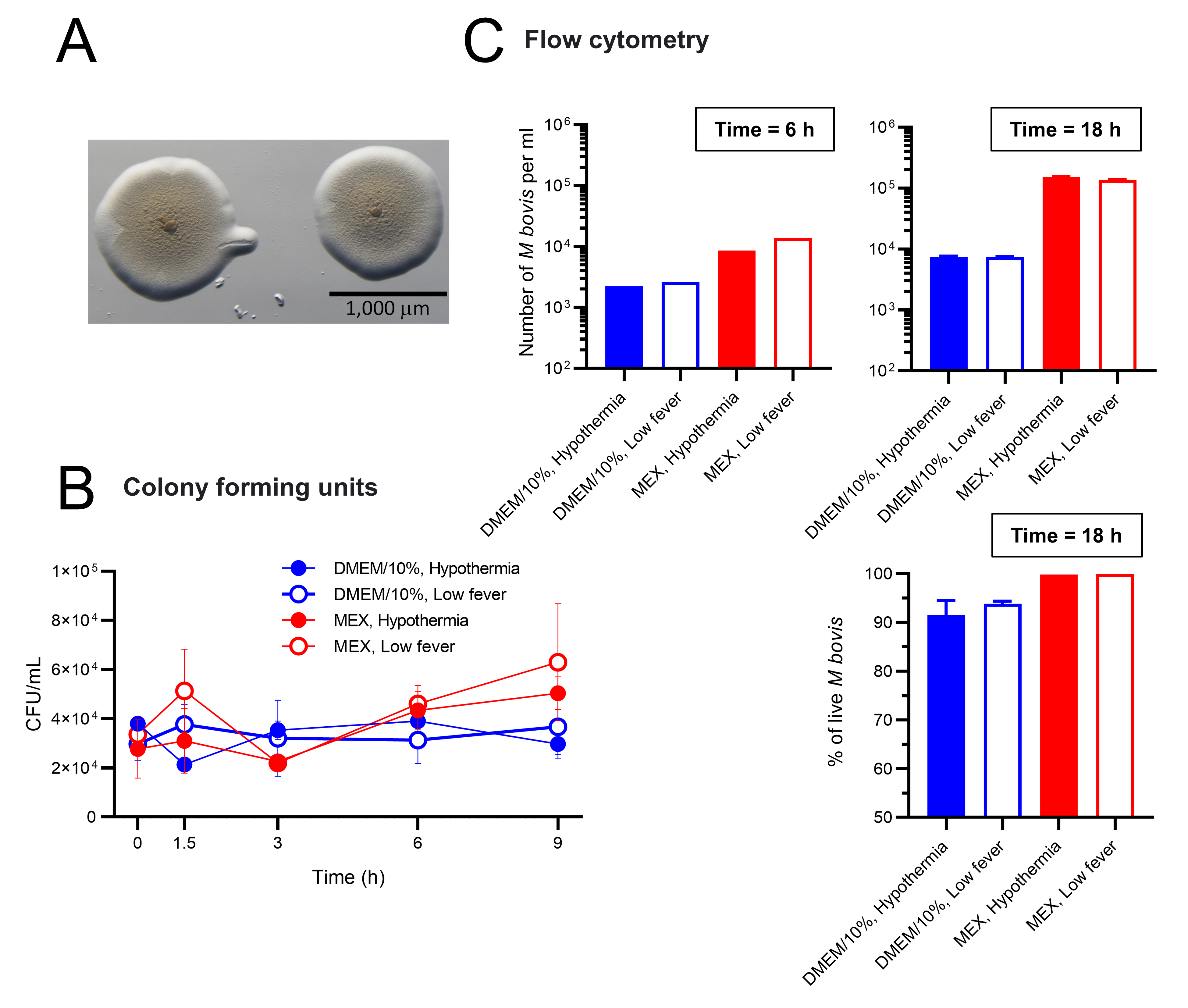

### TNF-alpha production in response to Mycoplasmopsis bovis is a concerted action of monocytes, B cells and NK cells

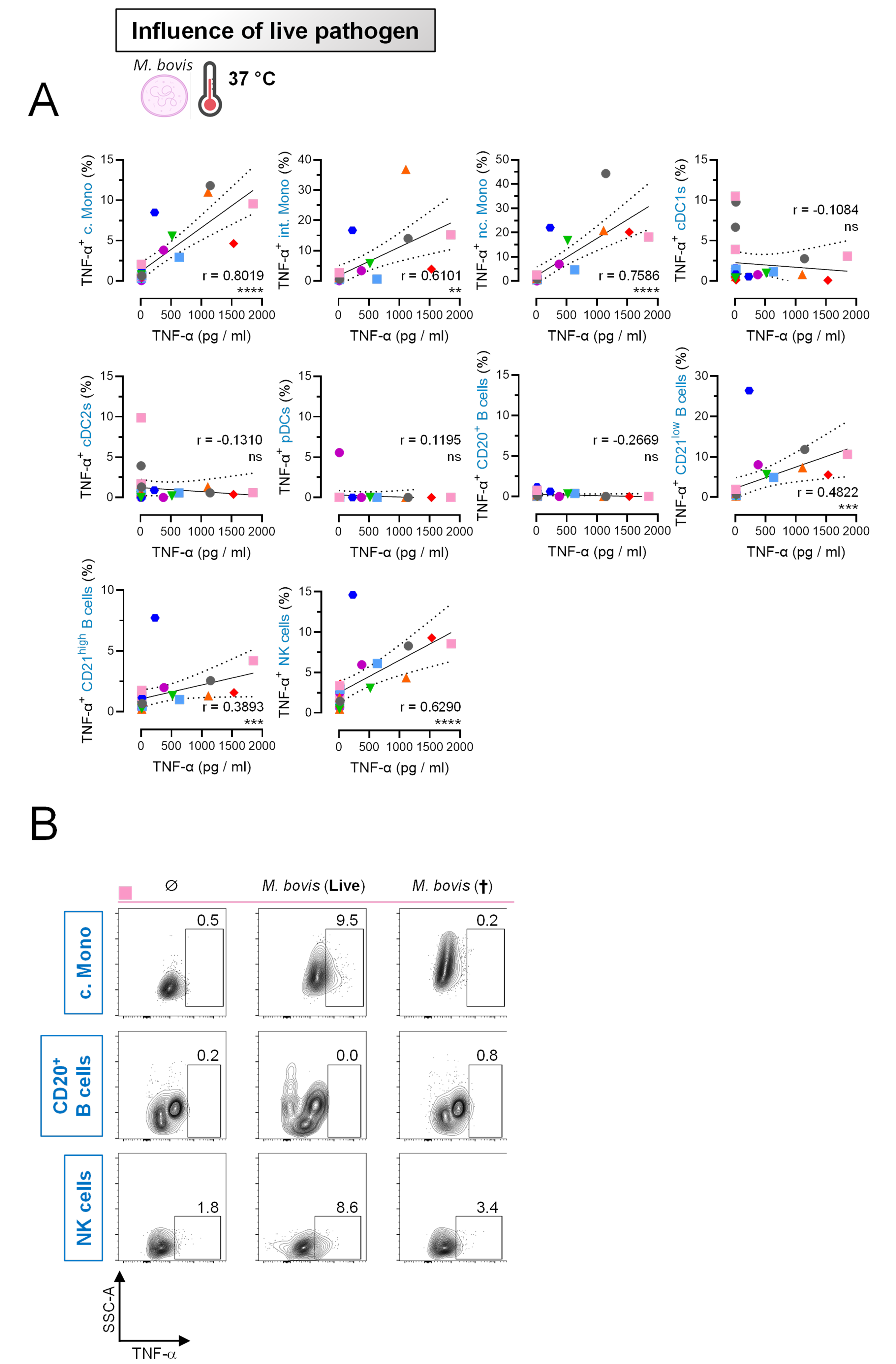
